# Improved Predictions of Phase Behaviour of Intrinsically Disordered Proteins by Tuning the Interaction Range

**DOI:** 10.1101/2022.07.09.499434

**Authors:** Giulio Tesei, Kresten Lindorff-Larsen

## Abstract

The formation and viscoelastic properties of condensates of intrinsically disordered proteins (IDPs) is dictated by amino acid sequence and solution conditions. Because of the involvement of biomolecular condensates in cell physiology and disease, furthering our understanding of the relationship between protein sequence and phase separation (PS) may have important implications in the formulation of new therapeutic hypotheses. Here, we present CALVADOS 2, a coarse-grained model of IDPs that accurately predicts conformational properties and propensity to undergo PS for diverse sequences and solution conditions. In particular, we systematically study the effect of varying the range of nonionic interactions and use our findings to improve the temperature scale of the model. We further optimize the residue-specific model parameters against experimental data on the conformational properties of 55 proteins, while also leveraging 70 hydrophobicity scales from the literature to avoid overfitting the training data. Extensive testing shows that the model accurately predicts chain compaction and PS propensity for sequences of diverse length and charge patterning, as well as at different temperatures and salt concentrations.

## 1 Introduction

Biomolecular condensates may form via phase separation (PS) into coexisting solvent-rich and macromolecule-rich phases. PS is driven by multiple transient interactions which are in many cases engendered by intrinsically disordered proteins (IDPs) and low-complexity domains (LCDs) of multi-domain proteins [1, 2, 3, 4, 5]. The propensity of IDPs to phase separate and the viscoelastic properties of the condensates are dictated by the amino acid sequence of the constituent IDPs. Moreover, condensates of some IDPs reconstituted in vitro tend to undergo a transition to a dynamically-arrested state, wherein oligomeric species can nucleate and ultimately aggregate into fibrils [6, 7, 2, 3, 8, 9, 10, 11, 12, 13, 14]. As accumulating evidence suggests that these processes may be involved in neurodegeneration and cancer [15, 16, 17, 18], understanding how PS and rheological properties of condensates depend on protein sequence is a current research focus. Due to the transient nature of the protein-protein interactions underpinning PS, quantitative characterization of biomolecular condensates via biophysical experimental methods is challenging, and hence molecular simulations have played an important role in aiding the interpretation of experimental data on condensates reconstituted in vitro [19]. Molecular simulations of the PS of IDPs require a minimal system size of ∼100 chains and long simulation times to sample the equilibrium properties of the two-phase system. Therefore, to enhance computational efficiency, it is useful to reduce the complexity of the system by modeling the solvent as a continuum while coarse-graining the atomistic representation of the protein to fewer interaction sites.

A widely used class of coarse-grained models of IDPs describes each residue as a single site centered at the C*α* atom. Charged residues interact via salt-screened electrostatic interactions whereas the remaining nonionic nonbonded interactions are incorporated in a single short-range potential characterized by a set of “stickiness” parameters. The “stickiness” parameters are specific to either the single amino acid or pairs of residues and were originally derived from a hydrophobicity scale [20]. Other models, based on the lattice simulation engines LaSSI [21] and PIMMS [22], classify the amino acids into a reduced number of residue types with distinct “stickiness”, ranging from binary categorizations into stickers and spacers [4, 23, 24] to more detailed descriptions and parameterizations [25, 26]. Recently, the accuracy of the “stickiness” parameters has been considerably improved. This has been achieved by leveraging (i) experimental data on single-chain properties, (ii) statistical analyses of protein structures, and (iii) residue-residue free energy profiles calculated from all-atom simulations [27, 28, 29, 30, 31, 26]. In particular, we and others have proposed an automated procedure to optimize the “stickiness” parameters so as to maximize the agreement with experimental small-angle X-ray scattering (SAXS) and paramagnetic relaxation enhancement (PRE) NMR data for a large set of IDPs [28, 29, 32, 26]. To ensure the transferability of the model across sequence space, we employed a Bayesian regularization approach [32, 33]. As the regularization term, we defined the prior knowledge on the “stickiness” parameters in terms of 87 hydrophobicity scales reported in the literature. The resulting M1 parameters capture the relative propensities to phase separate of a wide range of IDP sequences. However, we also observed that a systematic increase in simulation temperature of about 30 °C was needed to quantitatively reproduce the experimental concentration of the dilute phase coexisting with the condensate on an absolute scale. Herein, we refer to this model as the first version of the CALVADOS (Coarse-graining Approach to Liquid-liquid phase separation Via an Automated Data-driven Optimisation Scheme) model (CALVADOS 1).

In this class of coarse-grained models of IDPs, nonionic interactions are modeled via a Lennard-Jones-like potential, which decays to zero only at infinite residue-residue distances. For computational efficiency, the potential is typically calculated up to a cutoff distance, *r*_*c*_, and interactions between particles that are farther apart are ignored. Although this truncation may introduce severe artifacts, in the different implementations of the models, the value of *r*_*c*_ has varied considerably between 1 and 4 nm [29, 20, 34, 31, 30, 35, 32]. Here, we systematically investigate the effect of the cutoff of nonionic interactions on single-chain compaction and PS propensity. We find that decreasing the cutoff from 4 to 2 nm results in a small increase in the radius of gyration whereas the PS propensity significantly decreases. We exploit this effect to improve the temperature-dependence of the CALVADOS 1 model by tuning the cutoff of the nonionic potential. Further, we perform a Bayesian optimization of the “stickiness” parameters using a cutoff of 2.4 nm and an augmented training set. We show that the updated model (CALVADOS 2) has improved predictive accuracy.

## 2 Results and Discussion

When applying a cutoff scheme, we neglect the interactions of residues separated by a distance, *r*, larger than the cutoff, *r*_*c*_. For the most strongly interacting residue pair (between two tryptophans), the nonionic potential of the CALVADOS 1 model at *r*_*c*_ = 2 nm takes the value of -5 J mol^−1^, that is, only a small fraction of the thermal energy (Fig. 1*A*). However, the Lennard-Jones potential falls off slowly whereas the number of interacting partners increases quadratically with increasing *r*. Therefore, in a simulation of a protein-rich phase, decreasing the cutoff from 4 to 2 nm (Fig. 1*A*) can imply ignoring a total interaction energy per protein of several times the thermal energy.

**Figure 1:**
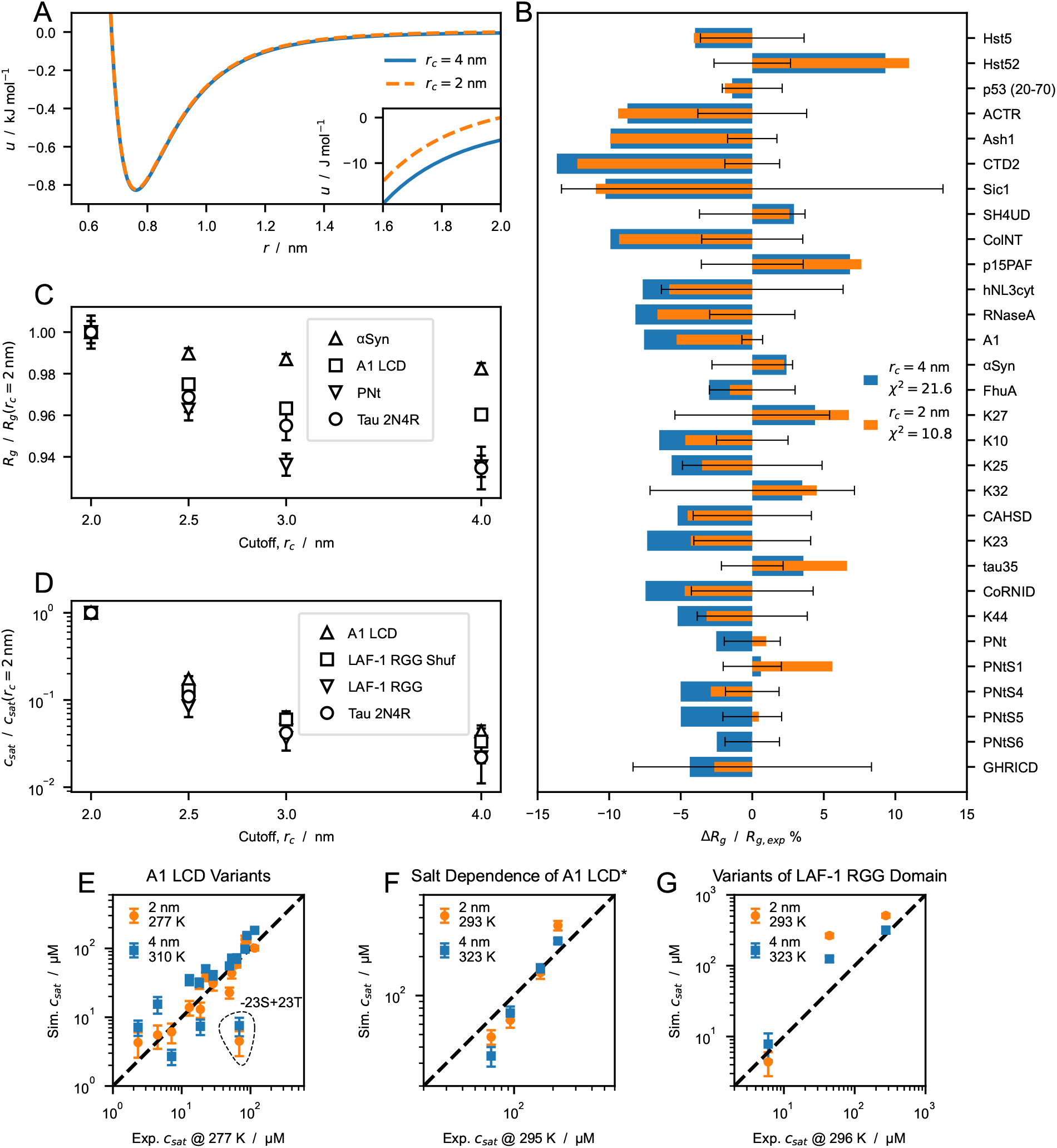
Effect of cutoff size on predictions of radii of gyration, *R*_*g*_, and saturation concentration, *c*_*sat*_, from simulations performed using the CALVADOS 1 parameters. (*A*) Nonionic Ashbaugh-Hatch potentials between two W residues with cutoff, *r*_*c*_, of 4 (blue solid line) and 2 nm (orange dashed line). The inset highlights differences between the potentials for *r*_*s*_ ≤ *r* ≤ *r*_*c*_. (*B*) Relative difference between experimental and predicted radii of gyration, ⟨*R*_*g*_⟩, for *r*_*c*_ = 4 (blue) and 2 nm (orange). 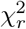 values reported in the legend are calculated for all the sequences in Table S1. (*C*) ⟨*R*_*g*_⟩ of *α*-Synuclein, hnRNPA1 LCD, PNt and human full-length tau (Table S1 and S2) from simulations performed with increasing cutoff size, *r*_*c*_, and normalized by the value at *r*_*c*_ = 2 nm. (*D*) Saturation concentration, *c*_*sat*_, for hnRNPA1 LCD, the randomly shuffled sequence of LAF-1 RGG domain, LAF-1 RGG domain and human full-length tau for increasing values of *r*_*c*_ and normalized by the *c*_*sat*_ at *r*_*c*_ = 2 nm. (*E*–*G*) Correlation between *c*_*sat*_ from simulations and experiments for (*E*) A1 LCD variants, (*F*) A1 LCD^∗^ WT at [NaCl] = 0.15, 0.2, 0.3 and 0.5 M and (*G*) variants of LAF-1 RGG domain (Table S4).

We first look into the effect of the choice of cutoff on the conformational ensembles of isolated proteins. We simulated single IDPs of different sequence length, *N* = 71–441, and average hydropathy, ⟨*λ* ⟩ = 0.33–0.63. The average radii of gyration, ⟨*R*_*g*_⟩, calculated from simulation trajectories are systematically larger when we use *r*_*c*_ = 2 nm, compared to the values obtained using *r*_*c*_ = 4 nm. CALVADOS 1 was optimized using the longer *r*_*c*_ and estimating the ensemble average*R*_*g*_ values as the root-mean-square 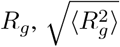. Since 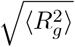 is systematically larger than ⟨*R*_*g*_⟩, decreasing *r*_*c*_ to 2 nm results in a slight improvement of the agreement between the calculated ⟨*R*_*g*_⟩ values and the experimental data (Fig. 1*B*).

To gain further insight into the effect of the cutoff, we performed simulations of single chains of *α*-Synuclein, hn-RNPA1 LCD, PNt and Tau 2N4R (Table S1 and S2) using *r*_*c*_ = 2, 2.5, 3 and 4 nm. Irrespective of the sequence, ⟨*R*_*g*_⟩ decreases monotonically with increasing *r*_*c*_. However, the effect on compaction appears to be larger for long sequences and high content of hydrophobic residues, both of which result in an increased number of shorter intramolecular distances. For example, upon increasing the *r*_*c*_ from 2 to 4 nm, the ⟨*R*_*g*_⟩ of *α*-Synuclein (*N* = 140, ⟨*λ*⟩ = 0.33) decreases by 2.3% whereas the effect is more pronounced for hnRNPA1 LCD (*N* = 137, ⟨*λ*⟩ = 0.61) and Tau 2N4R (*N* = 441, ⟨*λ*⟩ = 0.38), with a decrease in ⟨*R*_*g*_⟩ of 4.0% and 7.7%, respectively.

To investigate the effect of the cutoff distance on PS propensity, we performed direct-coexistence simulations of 100 chains of hnRNPA1 LCD, LAF-1 RGG domain (WT and shuffled sequence with higher charge segregation), and Tau 2N4R (Table S4). From the simulation trajectories of the two-phase system at equilibrium, we calculate *c*_*sat*_, i.e. the protein concentration in the dilute phase coexisting with the condensate. The higher the *c*_*sat*_ value, the lower the propensity of the IDP to undergo PS. As expected from the increased contact density in the condensed phase, the choice of cutoff has a considerably larger impact on *c*_*sat*_ than on chain compaction: decreasing *r*_*c*_ from 4 to 2 nm results in an increase in *c*_*sat*_ of over one order of magnitude. In contrast to what we observed for the ⟨*R*_*g*_⟩, the decrease in *c*_*sat*_ does not show a clear dependence on sequence length and average hydropathy. From the multi-chain trajectories of hnRNPA1 LCD, LAF-1 RGG domain (WT and shuffled sequence) and Tau 2N4R obtained using *r*_*c*_ = 4 nm, we estimate that the increase in nonionic energy per protein upon decreasing the cutoff from 4 to 2 nm is 13±1 kJ mol^−1^ (mean±standard deviation), respectively (Fig. S1*A*). Assuming that the number of interactions neglected by the shorter cutoff is proportional to the sequence length and the amino acid concentration in the condensate, the small variance in the energy increase across the different IDPs finds explanation in the fact that the simulated systems display similar values of *N* ^2^ × *c*_*con*_ (Fig. S1*A*), where *c*_*con*_ is the protein concentration in the condensate. The ratio *U* (*r*_*c*_ = 2 nm)*/U* (*r*_*c*_ = 4 nm) of the nonionic energies for *r*_*c*_ = 2 and 4 nm is also largely system independent (Fig. S1*B*). Moreover, decreasing the temperature by ∼30 K in the range between 310 and 323 K has a rather small impact on the relative strength of the electrostatic interactions with respect to the thermal energy, due to the decrease in the dielectric constant of water with increasing temperature (Fig. S1*C*). Therefore, we speculate that the effect of decreasing *r*_*c*_ can be compensated by simulating the system at a lower temperature (Fig. S1*B*).

With these considerations in mind, we use the CALVADOS 1 model with *r*_*c*_ = 2 nm to run direct-coexistence simulations of IDPs for which *c*_*sat*_ has been measured experimentally (Table S4), i.e. variants of hnRNPA1 LCD, hnRNPA1 LCD^∗^ at various salt concentrations, and LAF-1 RGG domain variants. As we have shown that decreasing the range of the nonionic interactions disfavours PS, we perform these simulations at the experimental temperatures, which are lower by ∼30 K than those required to reproduce the experimental *c*_*sat*_ values when the model is simulated with *r*_*c*_ = 4 nm (Fig. 1*E*–*G*). The two-fold decrease in *r*_*c*_ enables the model to quantitatively recapitulate the experimental *c*_*sat*_ data at the temperature at which the experiments were conducted. Notably, we show this for diverse sequences, across a wide range of ionic strengths, and for variants with different charge patterning and numbers of aromatic and charged residues. These results suggest that the range of interaction of the Lennard-Jones potential may be too large [36]. While the *r*^−6^ dependence is strictly correct for dispersion interactions between atoms, the nonionic potential of our model incorporates a variety of effective nonbonded interactions between residues, and hence the Lennard-Jones potential is not expected to capture the correct interaction range [31].

Since CALVADOS 1 was developed using *r*_*c*_ = 4 nm, we examined whether reoptimizing the model with the shorter cutoff could result in a comparably accurate model. As detailed in the Methods Section, we performed a Bayesian parameter-learning procedure [32] using an improved algorithm, an expanded training set (Table S1), and *r*_*c*_ = 2 nm. Fig. S2 shows that the new model tends to underestimate the *c*_*sat*_ values of the most PS-prone sequences. We hypothesize that during the optimization the reduction of attractive forces due to the shorter cutoff is overcompensated by an overall increase in *λ*. We tested this hypothesis by performing the optimization with increasing values of *r*_*c*_, in the range between 2.0 and 2.5 nm, and found that the *c*_*sat*_ values predicted from simulations performed with *r*_*c*_ = 2.0 nm increased monotonically with the *r*_*c*_ used for the optimization (Fig. S3). Using *r*_*c*_ = 2.4 nm for the optimization resulted in a model with improved accuracy compared to CALVADOS 1 (Fig. 2), especially for the PS of LAF-1 RGG domain and the −23S+23T variant of A1 LCD. To test the robustness of the approach, the optimization was carried out starting from *λ*_0_ = 0.5 for all the amino acids (Fig. 2) and from *λ*_0_ =M1 (Fig. S4 and S5). The difference between the resulting sets of optimal *λ* values (Fig. S5*A*) is lower than 0.08 for all the residues and exceeds 0.05 only for S, T and A. The model obtained starting from *λ*_0_ = 0.5 is more accurate at predicting PS propensities and will be referred to as CALVADOS 2 hereafter. The *λ* values of CALVADOS 1 and 2 differ mostly for K, T, A, M, and V, whereas the smaller deviations (|Δ*λ*| *<* 0.09) observed for Q, L, I, and F (Fig. 2*A*) are within the range of reproducibility of the method (Fig. S5*A*). Although CALVA-DOS 1 was optimized using *r*_*c*_ = 4 nm, predictions of single-chain compaction from simulations performed using *r*_*c*_ = 2 nm are more accurate for CALVADOS 1 than for CALVADOS 2. This result can be explained by the opposing effects of decreasing the cutoff and calculating *R*_*g*_ values as ⟨*R*_*g*_⟩ insteead of 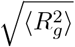. In fact, the 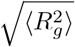 values predicted by CALVADOS 1 are strikingly similar to the ⟨*R*_*g*_⟩ values predicted by CALVADOS 2 (Fig. S6).

**Figure 2:**
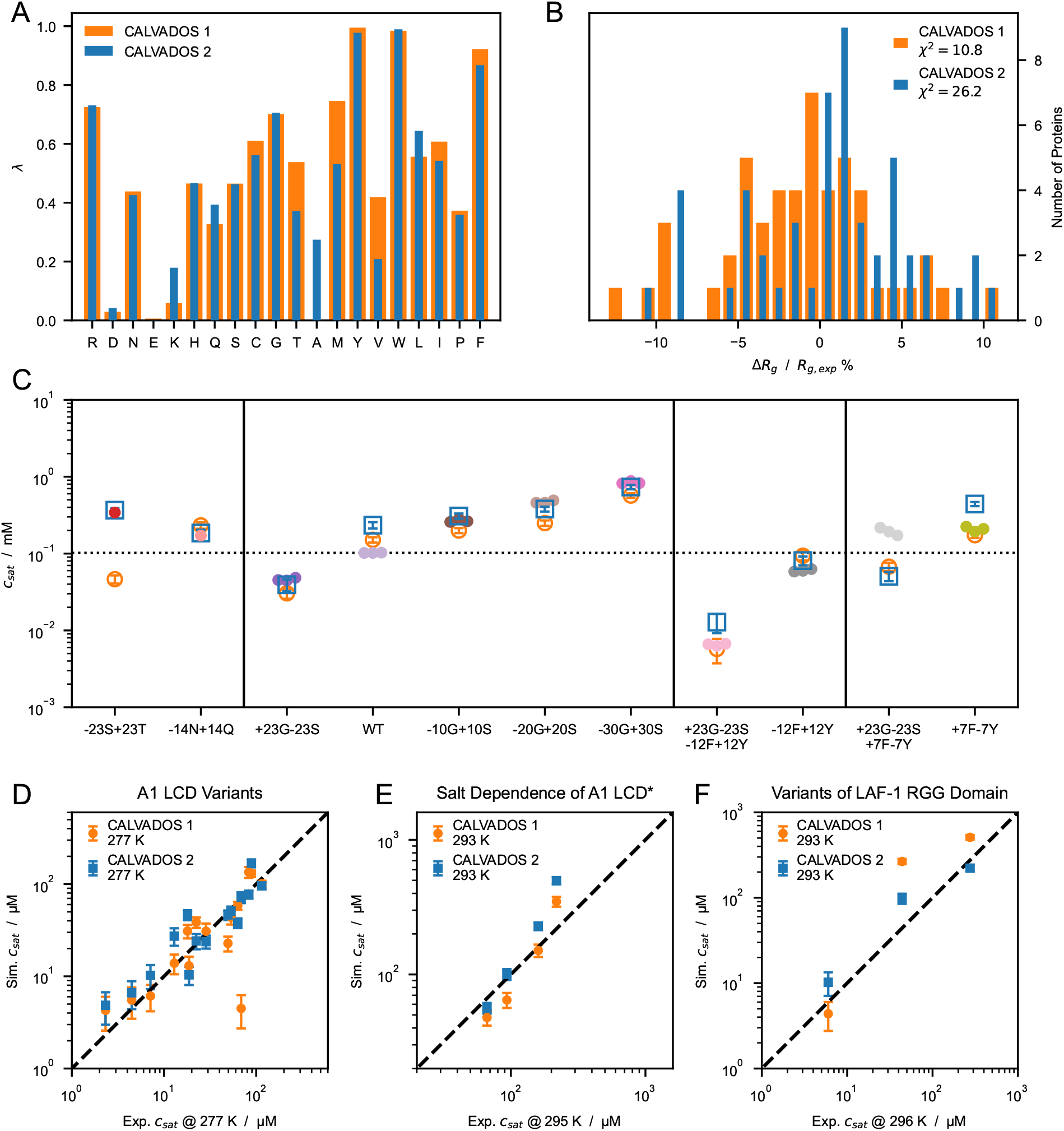
(*A*) Comparison between ***λ*** sets of CALVADOS 1 (orange) and CALVADOS 2 (blue). (*B*) Distribution of the relative difference between experimental (Table S1) and predicted radii of gyration, ⟨*R*_*g*_⟩, for CALVADOS 1 (orange) and CALVADOS 2 (blue). (*C*) Comparison between saturation concentrations, *c*_*sat*_, at 293 K of variants of hnRNPA1 LCD measured by Bremer, Farag, Borcherds et al. [37] (closed circles) and corresponding predictions of CALVADOS 1 (open orange circles) and CALVADOS 2 (open blue squares). (*D*–*F*) Correlation between *c*_*sat*_ from simulations and experiments for (*D*) A1 LCD variants, (*E*) A1 LCD^∗^ WT at [NaCl] = 0.15, 0.2, 0.3 and 0.5 M and (*F*) variants of LAF-1 RGG domain (Table S4).

The correlation between experimental and predicted *R*_*g*_ values for the 67 proteins of Table S1 and S2 is excellent for both CALVADOS 1 and 2 (Fig. S7*A*). On the other hand, CALVADOS 2 is more accurate than CALVADOS 1 at predicting PS propensities, as evidenced by Pearson’s correlation coefficients of 0.93 and 0.82, respectively, for the experimental and predicted *c*_*sat*_ values of the 26 sequences of Table S4 (Fig. S7*B*).

Capturing the interplay between short-range nonionic and long-range ionic interactions is essential for accurately modeling the PS of IDPs [38, 39, 37, 40]. Our results show that the decrease in the range of the nonionic potential reported in this work does not significantly perturb the balance between ionic and nonionic forces. In fact, CALVADOS 1 and 2 accurately predict the PS propensities of A1 LCD^∗^ at various salt concentrations, as well as the *c*_*sat*_ of variants of A1 LCD and LAF-1 RGG domain with different charge patterning (Fig. 1*E*–*G* and 2*D*–*F*). Moreover, CALVADOS 1 and 2 recapitulate the effect of salt concentration and charge patterning on the chain compaction of A1 LCD^∗^ [41] and p27-C constructs [42], respectively (Fig. S8).

As additional test systems, we considered constructs of the 1–80 N-terminal fragment of yeast Lge1, which have been recently investigated using turbidity measurements [43]. CALVADOS 2 correctly predicts that the WT Lge1_1−80_ construct undergoes PS at the experimental conditions, albeit with a hundred times larger *c*_*sat*_ (50 ± 6 µM at *c*_*s*_ = 100 mM) compared to experiments (*<* 1 µM). In agreement with experiments, CALVADOS 2 predicts that mutating all the 11 R residues to K increases *c*_*sat*_ by over one order of magnitude whereas mutating the 14 Y residues of the 1–80 fragment to A abrogates PS (Fig. S9).

## 3 Conclusions

In the context of a previously developed C*α*-based IDP model (CALVADOS), we show that neglecting the long range of attractive Lennard-Jones interactions has a small impact on the compaction of a single chain while strongly disfavouring PS. The effect can be explained by the smaller number of neglected pair interactions for a residues in an isolated chain compared to the dense environment of a condensate. Moreover, we find that the effect of reducing the range of interaction by a factor of two is relatively insensitive to sequence length and composition. Therefore, decreasing the cutoff of the Lennard-Jones potential of the C*α*-based model engenders a similar generic effect on chain compaction and PS as a corresponding increase in temperature. We take advantage of this finding to solve the temperature mismatch of the CALVADOS model. Namely, we decrease the cutoff of the nonionic interactions from 4 to 2 nm and obtain accurate *c*_*sat*_ predictions at the experimental conditions, whereas simulations at temperatures higher by 30 °C were required in the original implementation. Finally, we used the shorter cutoff to reoptimize the “stickiness” parameters of the model against experimental data reporting on single-chain compaction. The small expansion of the chain conformations is overcompensated by an overall increase in “stickiness” so that the resulting model tends to underestimate the experimental *c*_*sat*_ values. By systematically increasing the cutoff used in the development of the “stickiness” scale, we find that performing the optimization using *r*_*c*_ = 2.4 nm results in a model (CALVADOS 2) which yields accurate predictions from simulations run using *r*_*c*_ = 2 nm at the experimental conditions. We present CALVADOS 2 as an improvement of our previous model by testing on sets of experimental *R*_*g*_ and *c*_*sat*_ data comprising 16 and 36 systems, respectively, which were not used in the parameterization of the model.

## 4 Methods

### 4.1 Molecular Simulations

Molecular dynamics simulations are conducted in the NVT ensemble using the Langevin integrator with a time step of 10 fs and friction coefficient of 0.01 ps^−1^. Non-bonded interactions between residues separated by one bond are excluded from the energy calculations. Functional forms and parameters for bonded and nonbonded interactions are reported in the “Bonded and Nonbonded Interactions” Subsection. Single chains of *N* residues are simulated using HOOMD-blue v2.9.3 [44] in a cubic box of side length 0.38 × (*N* − 1) + 4 nm under periodic boundary conditions, starting from the fully extended linear conformation. Each chain is simulated in ten replicas for ∼ 6 × 0.3 × *N* ^2^ ps if *N >* 100 and for 18 ns otherwise. The initial 100 frames of each replica are discarded, so as to sample 5,000 weakly correlated conformations for each protein (Fig. S10). Direct-coexistence simulations are performed using openMM v7.5 [45] in a cuboidal box of side lengths [*L*_*x*_, *L*_*y*_, *L*_*z*_] = [25, 25, 300], [17, 17, 300] and [15, 15, 150] nm for Tau 2N4R, Ddx4 LCD, and for the remainder of the proteins, respectively. In the starting configuration, 100 chains are aligned along the *z*-axis and with their middle beads placed in the *xy*-plane at random (*x, y*) positions which are more than 0.7 nm apart. Multi-chain simulations are carried out for at least 2 µs, saving frames every 0.5 ns (Fig. S11, S12, and S13). After discarding the initial 0.6 µs, the slab is centered in the box at each frame as previously described [32] and the equilibrium density profile, *ρ*(*z*), is calculated by averaging over the trajectory of the system at equilibrium. The densities of the dilute and protein-rich phases are estimated as the average densities in the regions |*z*| *< z*_*DS*_ − *t/*2 and |*z*| *> z*_*DS*_ + 6*t* nm, where *z*_*DS*_ and *t* are the position of the dividing surface and the thickness of the interface, respectively. *z*_*DS*_ and *t* are obtained by fitting the semi-profiles in *z >* 0 and *z <* 0 to *ρ*(*z*) = (*ρ*_*a*_ + *ρ*_*b*_)*/*2 + (*ρ*_*b*_ − *ρ*_*a*_)*/*2 × tanh [(|*z*| − *z*_*DS*_)*/t*], where *ρ*_*a*_ and *ρ*_*b*_ are the densities of the protein-rich and dilute phases, respectively. The uncertainty of the density values is estimated as the standard error obtained from the blocking approach [46] implemented in the BLOCKING software (github.com/fpesceKU/BLOCKING).

### 4.2 Bonded and Nonbonded Interactions

In this study, we used the following truncated and shifted Ashbaugh-Hatch potential [47],

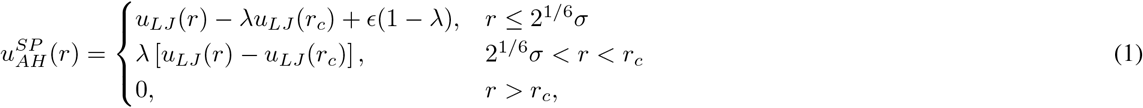

where *E* = 0.8368 kJ mol^−1^, *r*_*c*_ = 2 or 4 nm, and *u*_*LJ*_ is the Lennard-Jones potential:

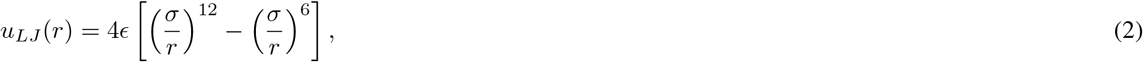

*σ* and *λ* are arithmetic averages of amino acid specific parameters quantifying size and hydropathy, respectively. For *σ*, we use the values calculated from van der Waals volumes by Kim and Hummer [48] whereas, for *λ*, we use the recently proposed M1 parameters [32] and the values optimized in this work.

Salt-screened electrostatic interactions are modeled via the Debye-Hückel potential,

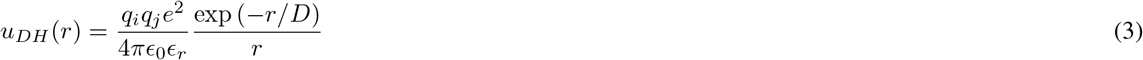

where *q* is the average amino acid charge number, *e* is the elementary charge, 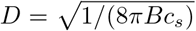 is the Debye length of an electrolyte solution of ionic strength *c*_*s*_ and *B*(ϵ_*r*_) is the Bjerrum length. Electrostatic interactions are truncated and shifted at the cutoff distance *r*_*c*_ = 4 nm, irrespective of the value of *r*_*c*_ used for Eq. 1. We use the following empirical relationship [49]

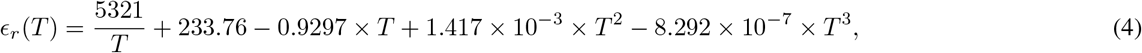

to model the temperature-dependent dielectric constant of the implicit aqueous solution. The Henderson–Hasselbalch equation is used to estimate the average charge of the histidine residues, assuming a p*K*_*a*_ value of 6 [50].

The amino acid beads are connected by harmonic potentials,

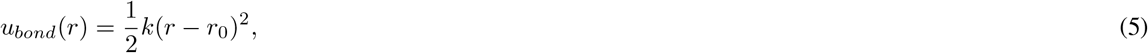

of force constant *k* = 8033 kJ mol^−1^ nm^−2^ and equilibrium distance *r*_0_ = 0.38 nm.

### 4.3 Optimization of the “Stickiness” Scale

The optimization of the “stickiness” parameters, ***λ***, is carried out to minimize the cost 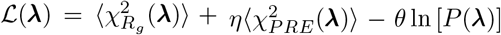 using an algorithm which is analogous to the one we previously described [32]. 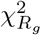 and 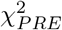 quantify the discrepancy between model predictions and experimental data, and are defined as 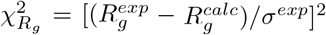 and 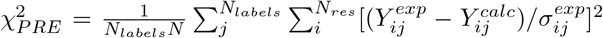, respectively, where *σ*^*exp*^ is the error on the experimental values, *Y* is either PRE rates or intensity ratios and *N*_*labels*_ is the number of spin-labeled mutants used for the NMR PRE data. In the expression for the cost function, the coefficients are set to *η* = 0.1 and *θ* = 0.05. The prior is the distribution of *λ, P* (***λ***), derived from a subset of the hydrophobicity scales reported in Table 3 and 4 of Simm et al. [51]. Specifically, only the 70 scales that are unique after min-max normalization (Fig. S14) are used for the calculation of *P* (*λ*), namely Wimley, BULDG reverse, MANP780101, VHEG790101, JANIN, JANJ790102, WOLR790101, PONP800101–6, Wilson, FAUCH, ENGEL, ROSEM, JACWH, CowanWhittacker, ROSM880101 reverse, ROSM880102 reverse, COWR900101, BLAS910101, CASSI, CIDH920101, CIDH920105, CIDBB, CIDA+, CIDAB, PONG1–3, WILM950101–2, WILM950104, Bishop reverse, NADH010101–7, ZIMJ680101, NOZY710101, JONES, LEVIT, KYTJ820101, SWER830101, SWEET, EISEN, ROSEF, GUYFE, COHEN, NNEIG, MDK0, MDK1, JURD980101, SET1–3, CHOTA, CHOTH, Sweet & Eisenberg, KIDER, ROSEB, Welling reverse, Rao & Argos, GIBRA, and WOLR810101 reverse. *P* (***λ***) is obtained via multivariate kernel density estimation, as implemented in scikit-learn [52], using a Gaussian kernel with bandwidth of 0.05. This prior is 20-dimensional and contains information on the *λ*-distribution of the single amino acid (Fig. S15) as well as on the covariance matrix (Fig. S16) inferred from our selection of 70 hydrophobicity scales.

In the first step of the optimization procedure, the *λ* values for all the amino acids are set to 0.5, *λ*_0_ = 0.5, and these parameters are used to simulate the proteins of the training set (Table S1). We proceed with the first optimization cycle, wherein, at each *k*-th iteration, the *λ* values of a random selection of five amino acids are nudged by random numbers picked from a normal distribution of standard deviation 0.05 to generate a trial ***λ***_***k***_ set. For each *i*th frame, we calculate the Boltzmann weight as *w*_*i*_ = exp {−[*U* (***r***_***i***_, ***λ***_***k***_) − *U* (***r***_***i***_, ***λ***_**0**_)]*/k*_*B*_*T* }, where *U* is the nonionic potential. The trial ***λ***_***k***_ is discarded if the effective fraction of frames, 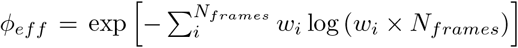, is lower than 60%. Otherwise, the acceptance probability follows the Metropolis criterion, min 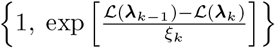, where *ξ*_*k*_ is a unitless control parameter. Each optimization cycle is divided into ten micro-cycles, wherein the control parameter, *ξ*, is initially set to *ξ*_0_ = 0.1 and scaled down by 1% at each iteration until *ξ <* 10^−8^. From the complete optimization cycle, we select the ***λ*** set yielding the lowest estimate of L. Consecutive optimization cycles are performed from simulations runs carried out with the intermediate optimal ***λ*** set. To show that the procedure is reproducible and that the final ***λ*** set is relatively independent of the initial conditions, we performed an additional optimization procedure starting from the M1 model, *λ*_0_ =M1 [32]. The optimization performed in this work differs from our previous implementation [32] also for the following details: (i) nine additional sequences have been included in the training set (Table S1 and S3); (ii) single chains are simulated as detailed in the “Molecular Simulations” Subsection; (v) the average radius of gyration is calculated as ⟨*R*_*g*_⟩ instead of 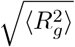.

## Code and Data Availability

Scripts and parameters to perform single-chain and direct-coexistence simulations using CALVADOS 1 and 2 are available at github.com/KULL-Centre/CALVADOS and archived on Zenodo at doi.org/10.5281/zenodo.6815068. All code and data to reproduce the results presented in this work are available at github.com/KULL-Centre/papers/tree/main/2022/CG-cutoffs-Tesei-et-al.

## Acknowledgments

We thank Anna Ida Trolle for her help in setting up the protocol for single-chain simulations. We thank Rosana Collepardo, Jerelle A. Joseph, and Aleks Reinhardt for useful discussions that led us to explore the effects of cutoffs. We acknowledge access to computational resources from the ROBUST Resource for Biomolecular Simulations (supported by the Novo Nordisk Foundation; NNF18OC0032608) and Biocomputing Core Facility at the Department of Biology, University of Copenhagen. This project has received funding from the European Union’s Horizon 2020 research and innovation programme under the Marie Skłodowska-Curie grant agreement No. 101025063.

## Supporting Figures and Tables

**Figure S1:**
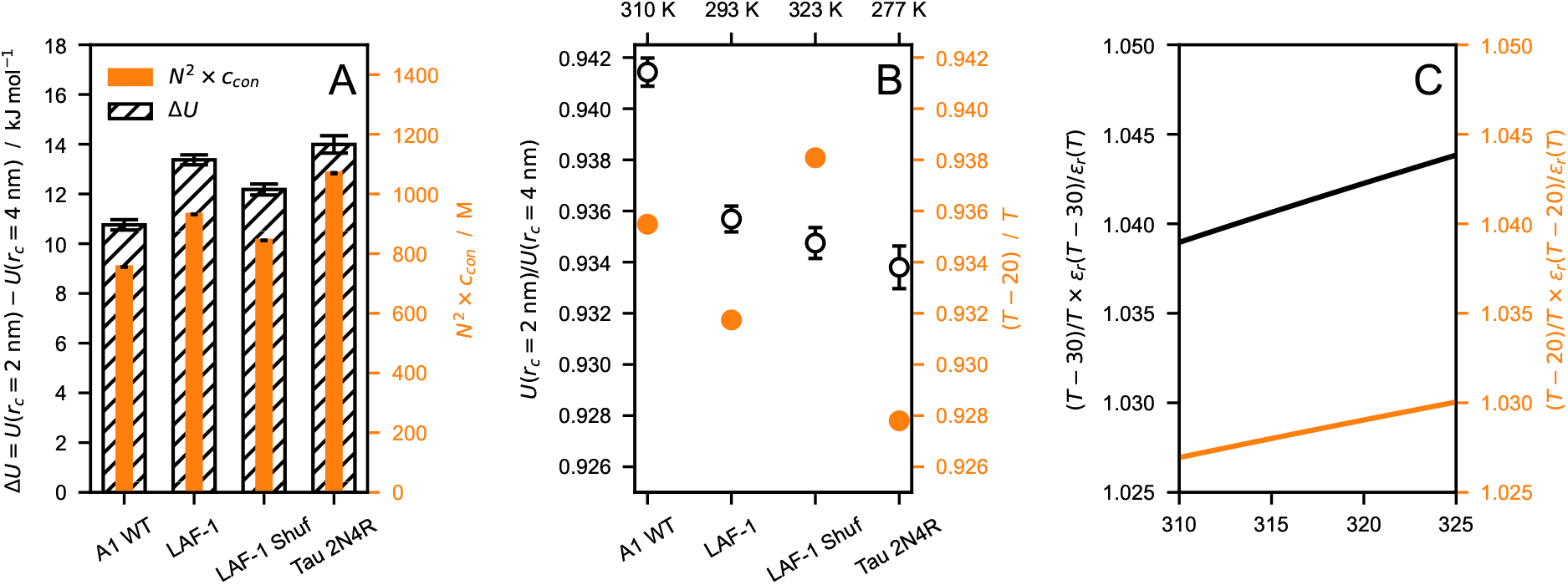
(*A*) Comparison between nonionic energy difference per protein (Δ*U* = *U* (*r*_*c*_ = 2 nm) − *U* (*r*_*c*_ = 4 nm), hatched) and *N* ^2^ × *c*_*con*_ (orange), where *N* is the sequence length and *c*_*con*_ is the molar protein concentration in the condensate. (*B*) Ratio between nonionic energies calculated with *r*_*c*_ = 2 and 4 nm (open circles) compared to the ratio of the thermal energy at *T*^′^ = *T* − 20 K and at *T* (orange). (*C*) Increase in electrostatic energy relative to the thermal energy upon decreasing the temperature by 30 (black) and 20 K (orange). The data shown in this figure are obtained from simulations of hnRNPA1 LCD, LAF-1 RGG domain (WT and shuffled sequence) and Tau 2N4R performed at *T* = 310, 293, 323, and 277 K, respectively, and using *r*_*c*_ = 4 nm. Error bars are standard deviations over trajectories of the systems at equilibrium.

**Figure S2:**
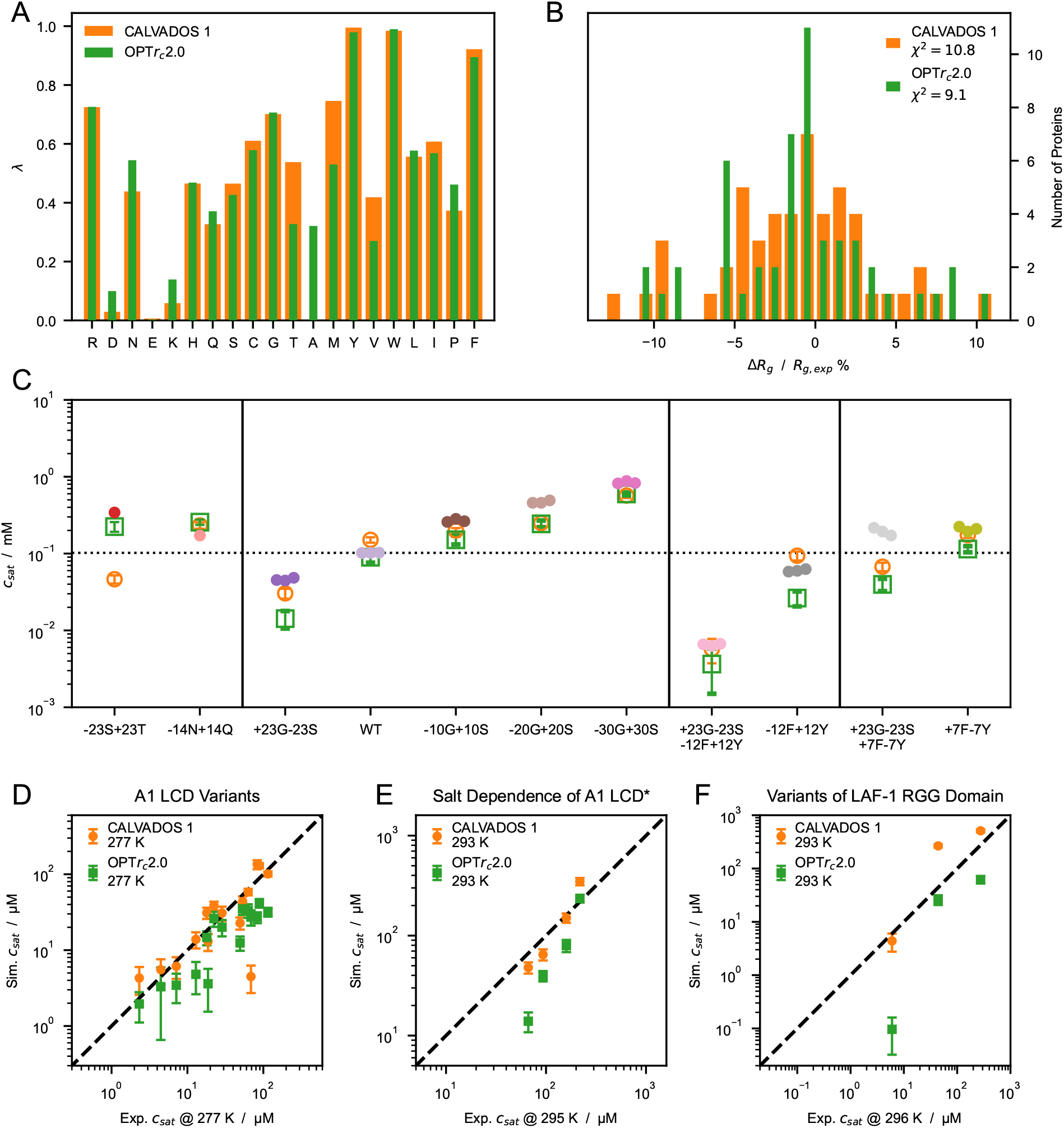
(*A*) Comparison between ***λ*** sets of CALVADOS 1 (orange) and the model resulting from the optimization with *r*_*c*_ = 2.0 nm (OPT*r*_*c*_2.0, green). (*B*) Distribution of the relative difference between experimental (Table S1) and predicted radii of gyration, ⟨*R*_*g*_⟩, for CALVADOS 1 (orange) and OPT*r*_*c*_2.0 (blue). (*C*) Comparison between saturation concentrations, *c*_*sat*_, at 293 K of variants of hnRNPA1 LCD measured by Bremer, Farag, Borcherds et al. [37] (closed circles) and corresponding predictions of CALVADOS 1 (open orange circles) and OPT*r*_*c*_2.0 (open green squares). (*D*–*F*) Correlation between *c*_*sat*_ from simulations and experiments for (*D*) A1 LCD variants, (*E*) A1 LCD^∗^ WT at [NaCl] = 0.15, 0.2, 0.3 and 0.5 M and (*F*) variants of LAF-1 RGG domain (Table S4).

**Figure S3:**
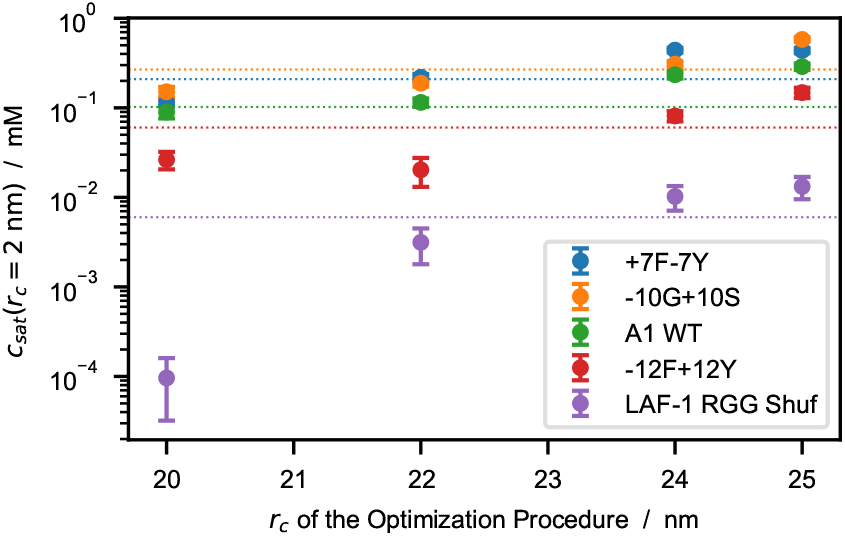
Saturation concentrations, *c*_*sat*_, as a function of the cutoff used to optimize the model. *c*_*sat*_ values are calculated from simulations performed using *r*_*c*_ = 2.0 nm whereas the models are optimized using *r*_*c*_ = 2.0, 2.2, 2.4, and 2.5 nm. Horizontal dotted lines represent experimental *c*_*sat*_ values from the references reported in Table S4.

**Figure S4:**
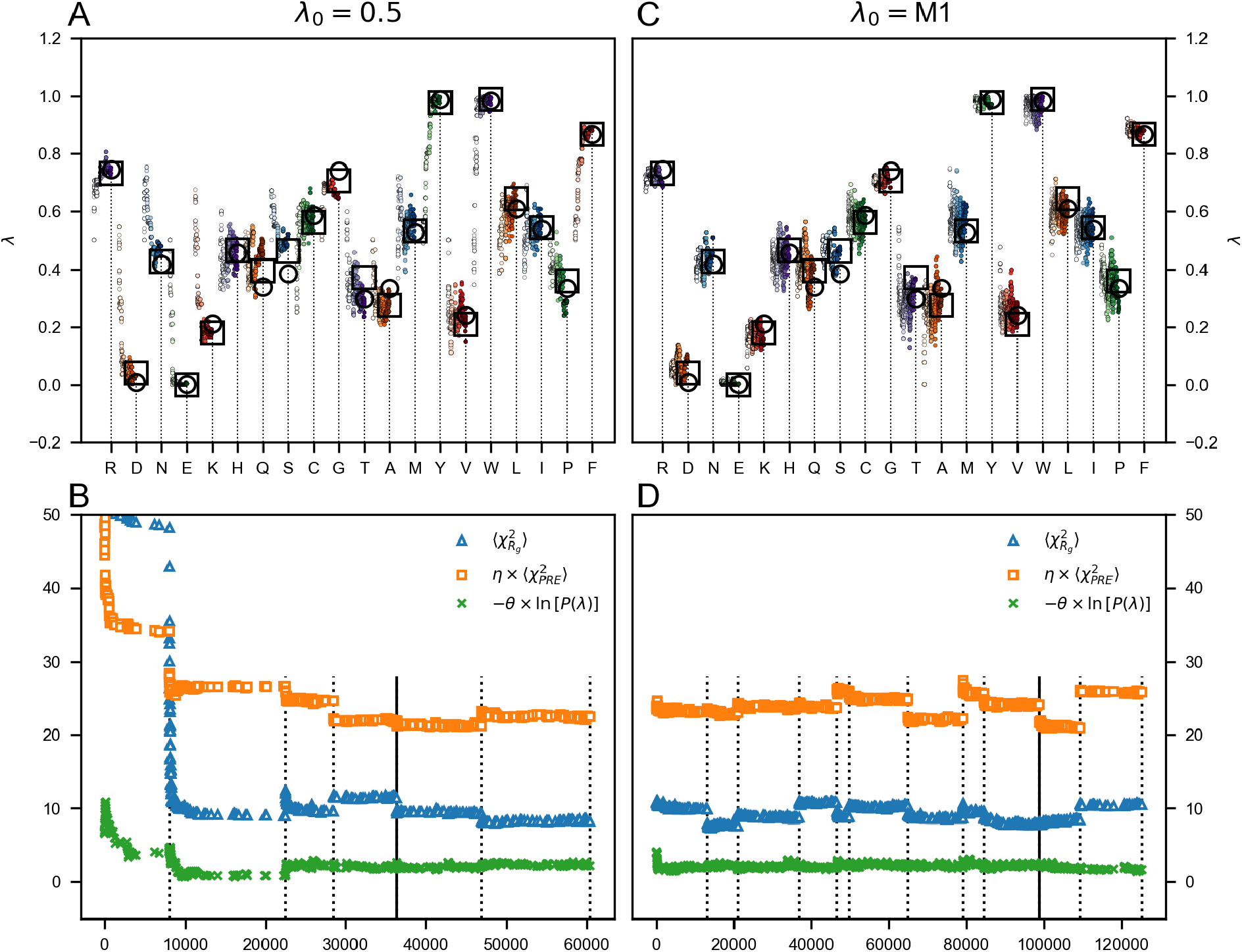
Optimization of the *λ* parameters starting from *λ*_0_ = 0.5 (*A* and *B*) and *λ*_0_ =M1 (*C* and *D*). (*A* and *B*) Evolution of the *λ* parameters during three consecutive optimization cycles. The color gradient from light to dark shade indicates increasing number of iterations. Open squares and circles show optimal *λ* sets obtained from indipendent optimizations starting from *λ*_0_ = 0.5 and *λ*_0_ =M1, respectively. (*C* and *D*) Evolution of 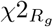 (blue triangles), 0.1 × *χ*2_*PRE*_ (orange squares), and the regularization term 0.05 × ln [*P* (***λ***)] (green circles). Dotted vertical lines indicate updated sampling by molecular simulations, whereas the remaining points are estimated from reweighted ensembles. Solid vertical lines indicate the optimal *λ* set corresponding to the lowest total cost function, ℒ.

**Figure S5:**
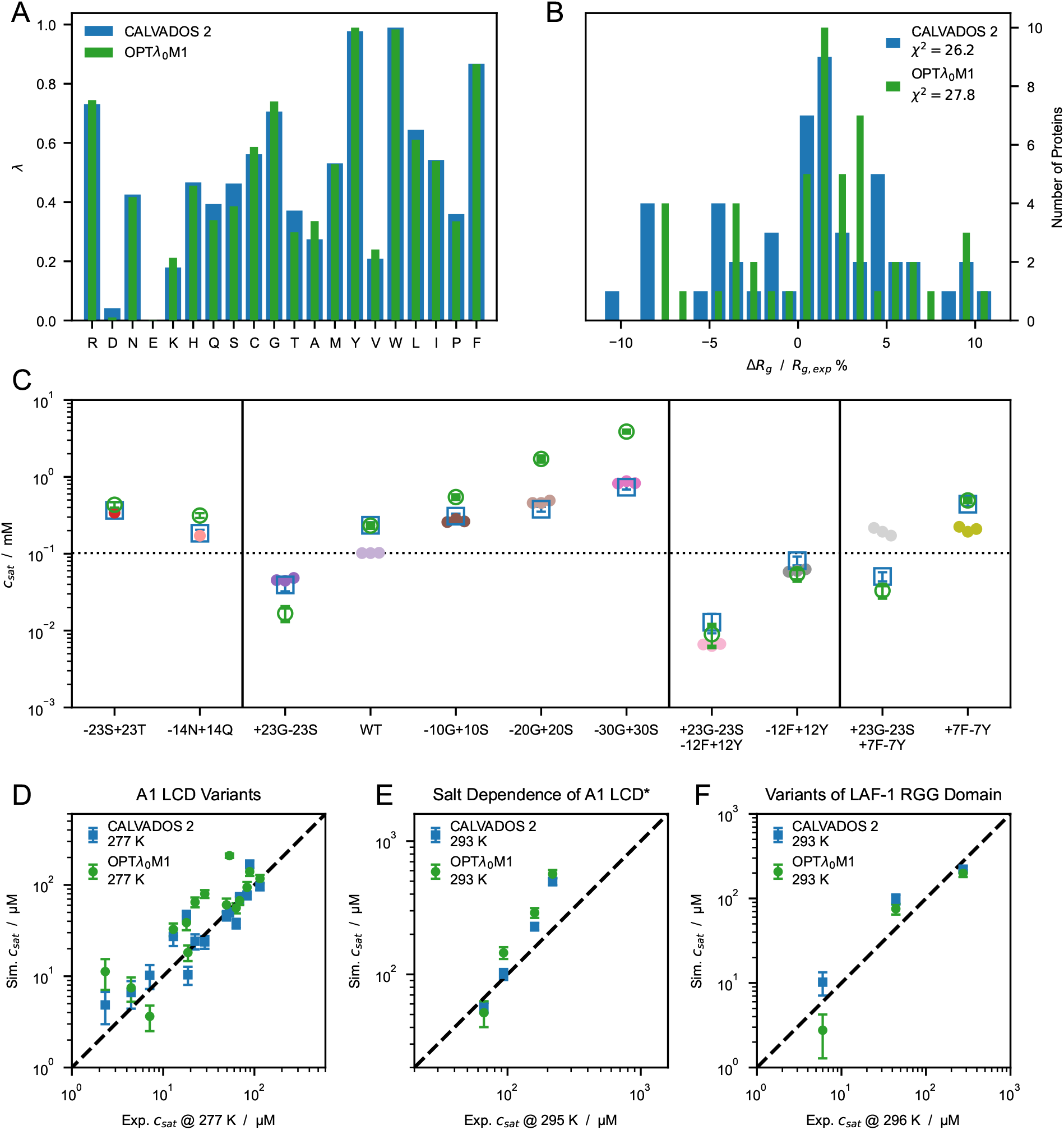
(*A*) Comparison between ***λ*** sets optimized starting from *λ*_0_ = 0.5 (CALVADOS 2, blue) and *λ*_0_ =M1 (OPT*λ*_0_M1, green) using *r*_*c*_ = 2.4 nm. (*B*) Distribution of the relative difference between experimental (Table S1) and predicted radii of gyration, ⟨*R*_*g*_⟩, for CALVADOS 2 (blue) and OPT*λ*_0_M1 (green). (*C*) Comparison between saturation concentrations, *c*_*sat*_, at 293 K of variants of hnRNPA1 LCD measured by Bremer, Farag, Borcherds et al. [37] (closed circles) and corresponding predictions of CALVADOS 2 (open blue squares) and OPT*λ*_0_M1 (open green circles). (*D*–*F*) Correlation between *c*_*sat*_ from simulations and experiments for (*D*) A1 LCD variants, (*E*) A1 LCD^∗^ WT at [NaCl] = 0.15, 0.2, 0.3 and 0.5 M and (*F*) variants of LAF-1 RGG domain (Table S4).

**Figure S6:**
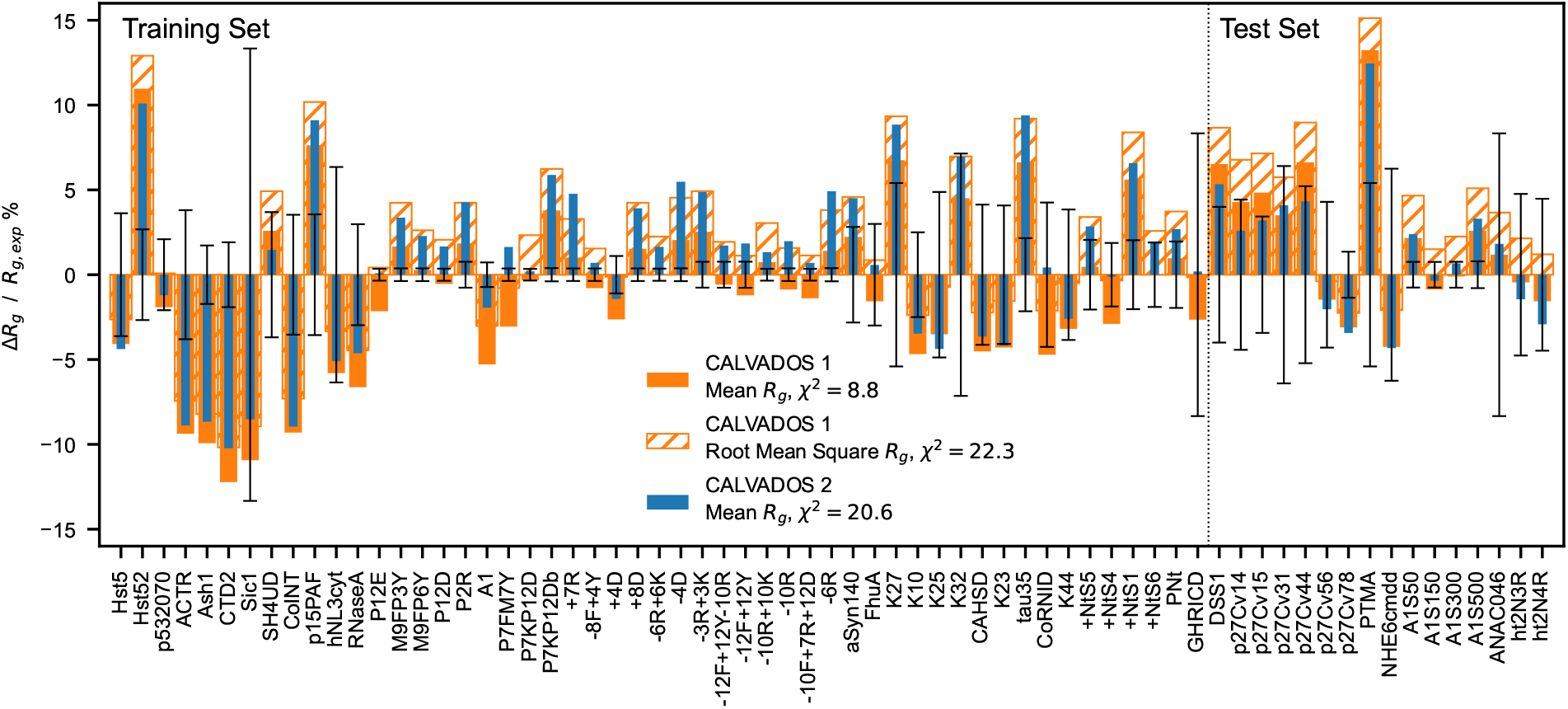
Relative difference between experimental and predicted radii of gyration for CALVADOS 1 (orange) and CALVADOS 2 (blue). Full and hatched bars show 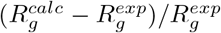 where 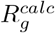 is calculated as the mean 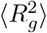 or the root mean square 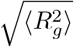, respectively. The vertical dashed line split the plot into the 51 sequences of the training set (Table S1) and the 16 sequences of the test set (Table S2).

**Figure S7:**
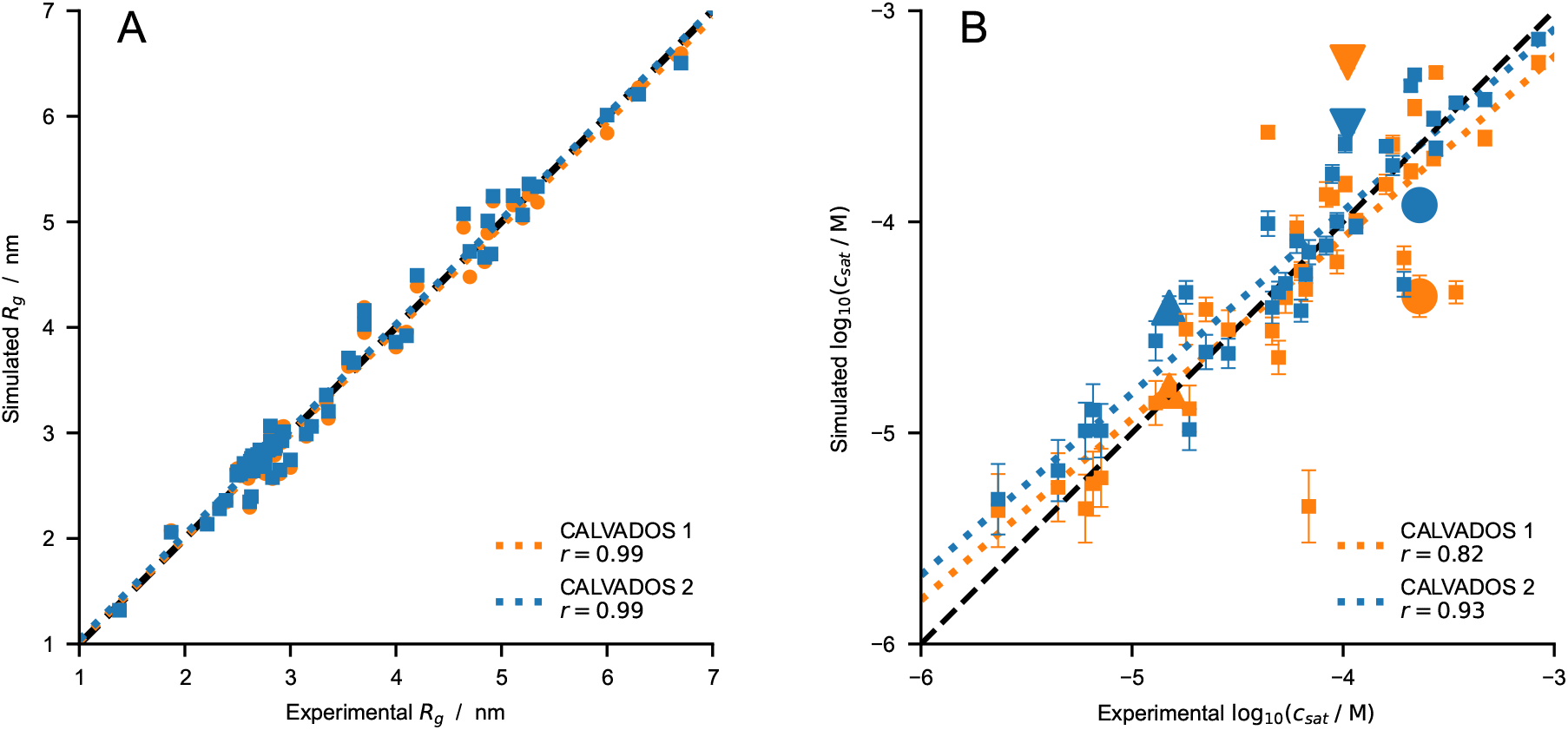
Comparison between experimental and predicted (*A*) *R*_*g*_ (Table S1 and S2) and (*B*) *c*_*sat*_ values for CALVADOS 1 (orange) and CALVADOS 2 (blue). Pearson’s *r* coefficients are reported in the legend. Small squares in *B* show the same data as in Fig. 2*C*–*F* whereas the large upward triangle, downward triangle, and circle show values for A2 LCD, FUS LCD, and Ddx4 LCD, respectively, at the conditions reported in Table S4.

**Figure S8:**
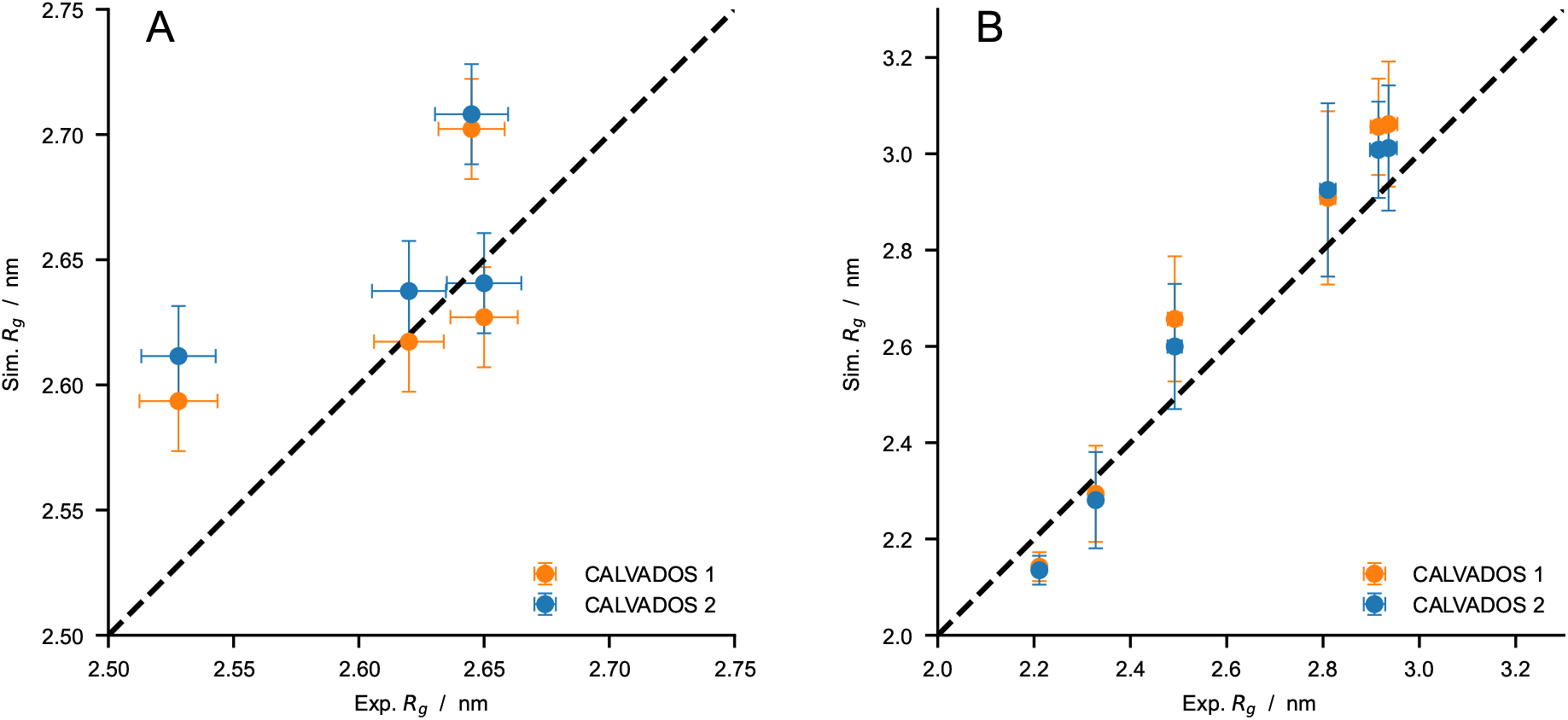
Comparison between experimental *R*_*g*_ values and predictions of CALVADOS 1 (orange) and CALVADOS 2 (blue) for (*A*) A1 LCD^∗^ at different salt concentrations (50 mM *< c*_*s*_ *<* 500 mM) and (*B*) p27-C constructs of different charge patterning (0.1 *< κ <* 0.8). Experimental conditions and references are reported in Table S2.

**Figure S9:**
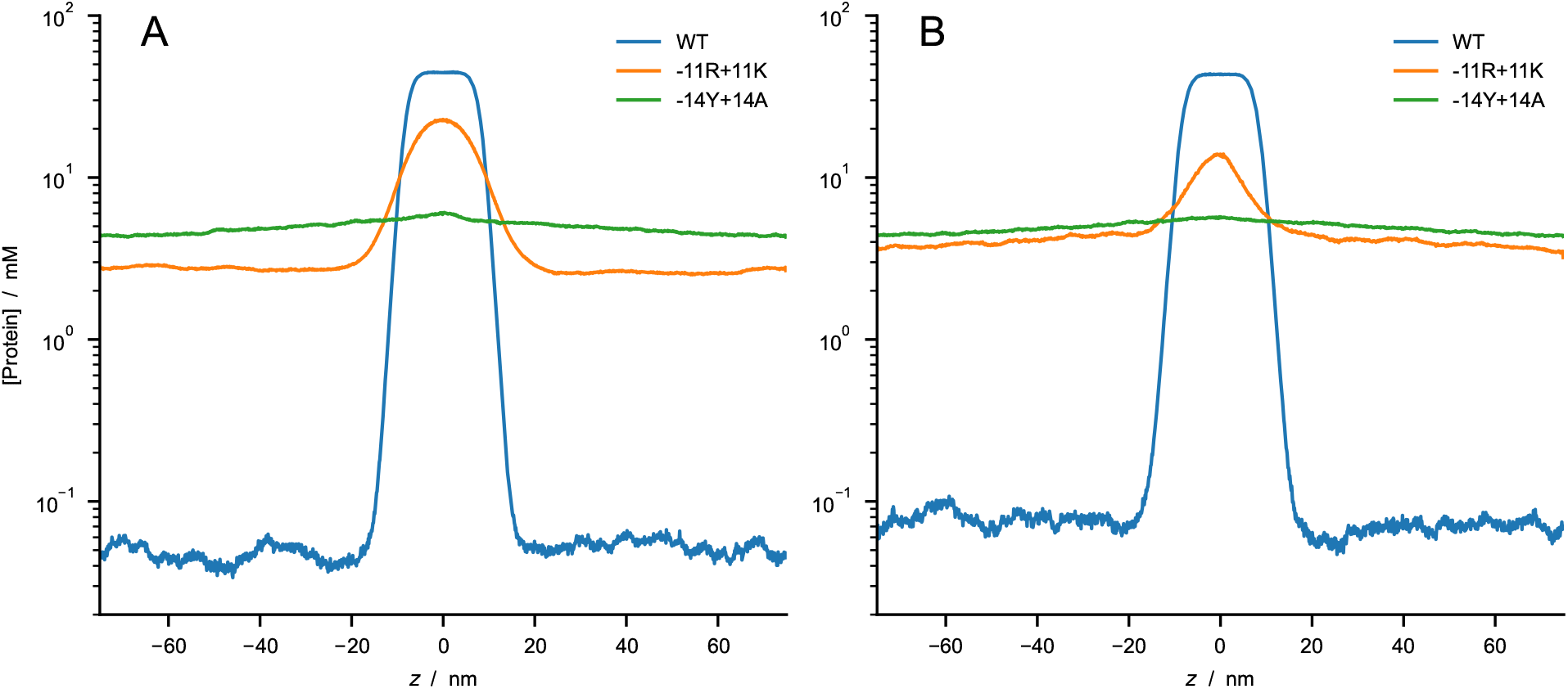
Predictions of CALVADOS 2 direct-coexistence simulations of the PS of constructs of the 1–80 N-terminal fragment of yeast Lge1 simulated at (*A*) *c*_*s*_ = 100 mM and (*B*) 500 mM. Protein concentration profiles are shown as a function of the long side of the simulation cell for WT (blue), -11R+11K variant (orange), and -14Y+14A variant (green).

**Figure S10:**
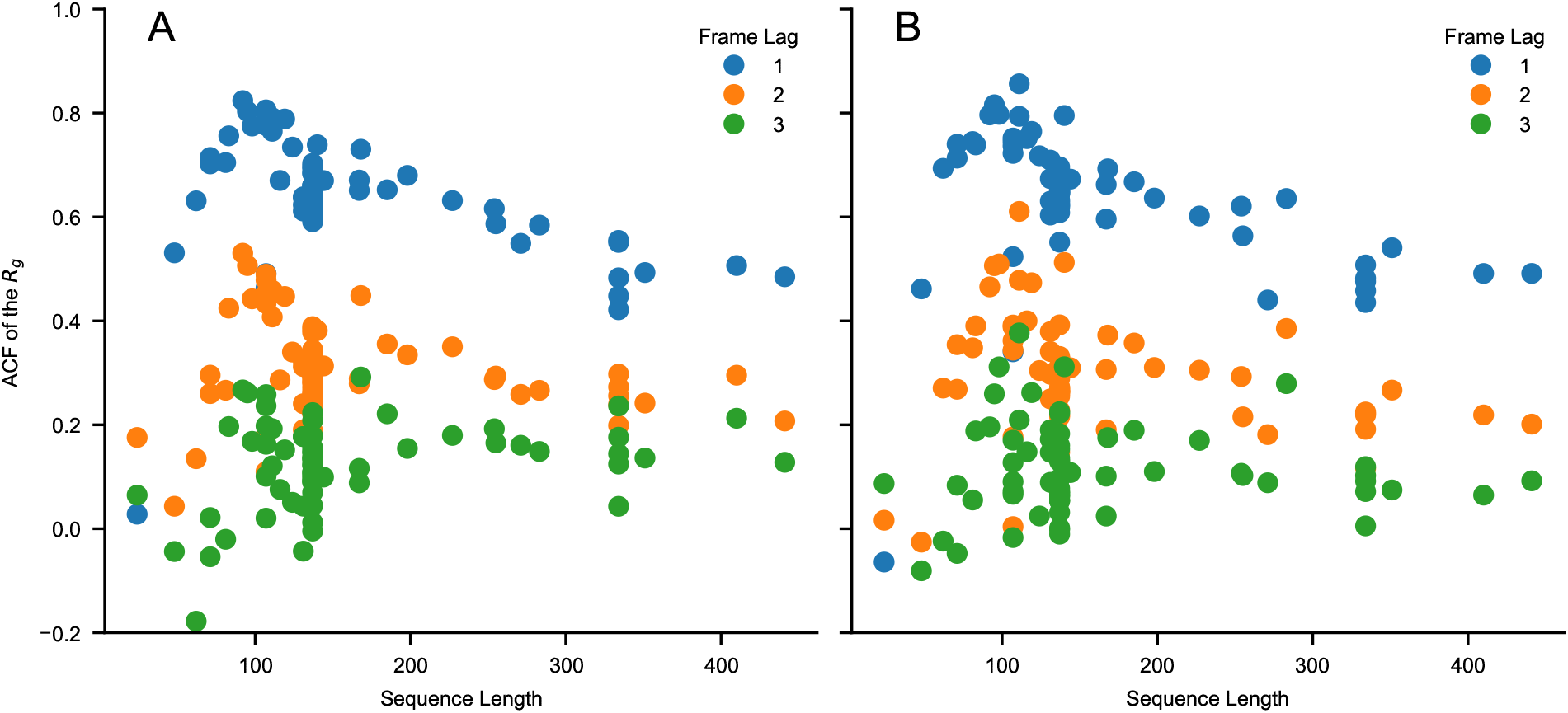
Values of the autocorrelation function of the *R*_*g*_ for a lag time of one, two and three frames as a function of sequence length, *N*. The autocorrelation is calculated for the sequences of Table S1 and S2 simulated using (*A*) CALVADOS 1 and (*B*) CALVADOS 2 for ∼ 6 × 0.3 × *N* ^2^ ps if *N >* 100 and for 18 ns otherwise. 600 simulation frames are saved every ∼ 0.003 × *N* ^2^ ps if *N >* 100 and every 30 ps otherwise. The initial 100 frames are discarded from each simulation.

**Figure S11:**
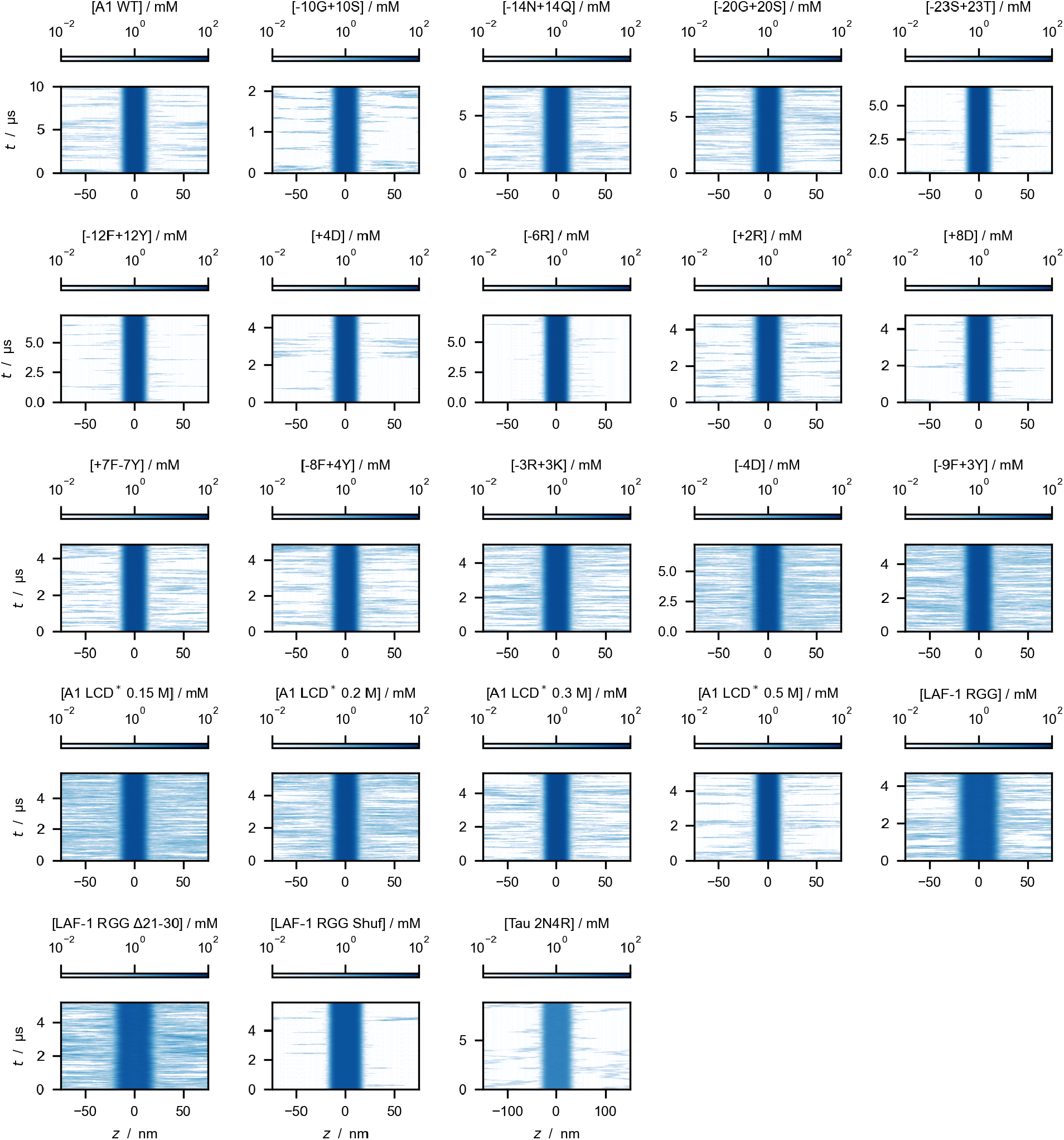
Time evolution of the protein concentration along the *z*-axis of the simulation cell, as obtained from direct-coexistence simulations performed with the CALVADOS 1 model and *r*_*c*_ = 4 nm.

**Figure S12:**
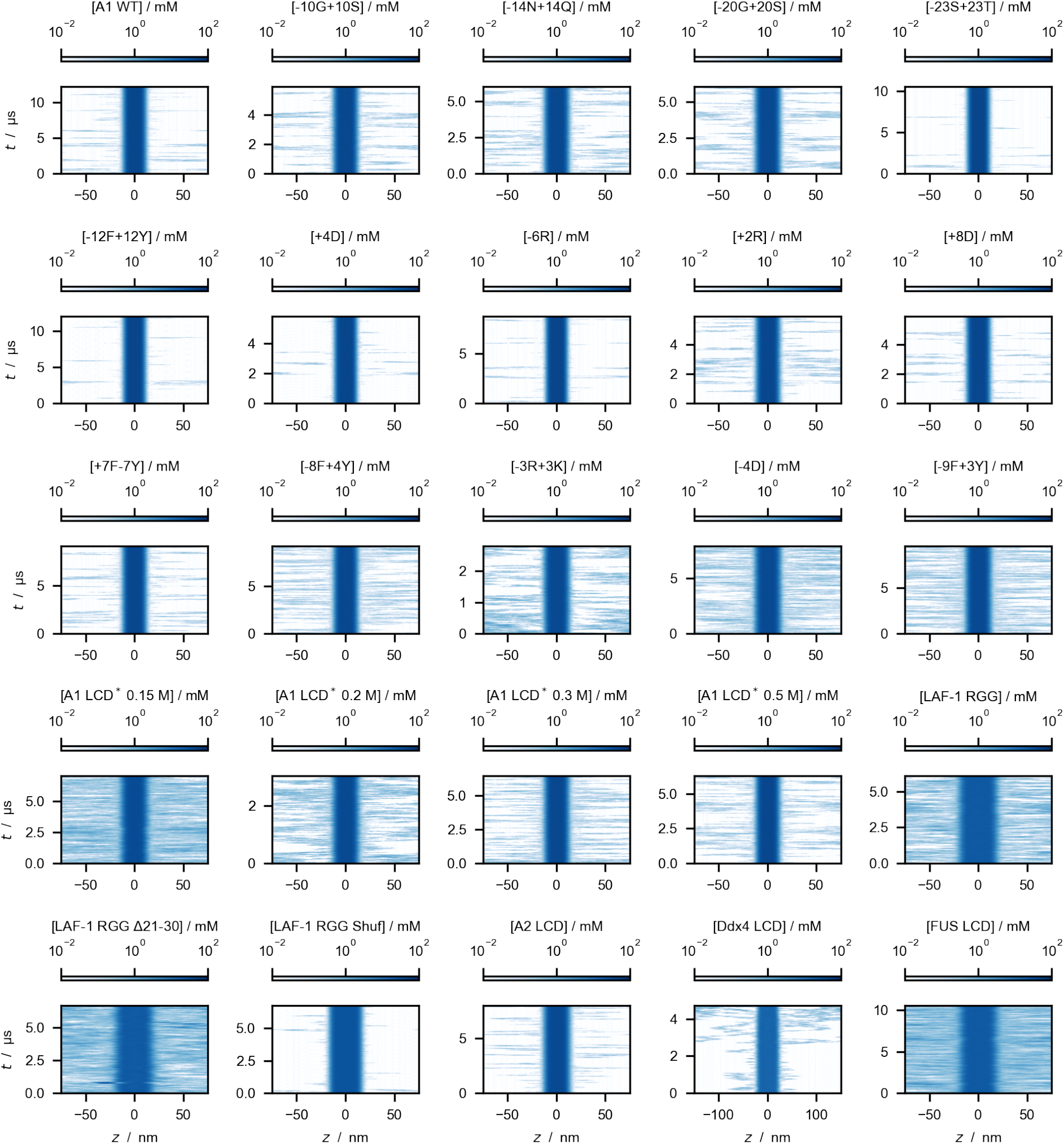
Time evolution of protein density along the *z*-axis of the simulation cell, as obtained from direct-coexistence simulations performed with the CALVADOS 1 model and *r*_*c*_ = 2 nm.

**Figure S13:**
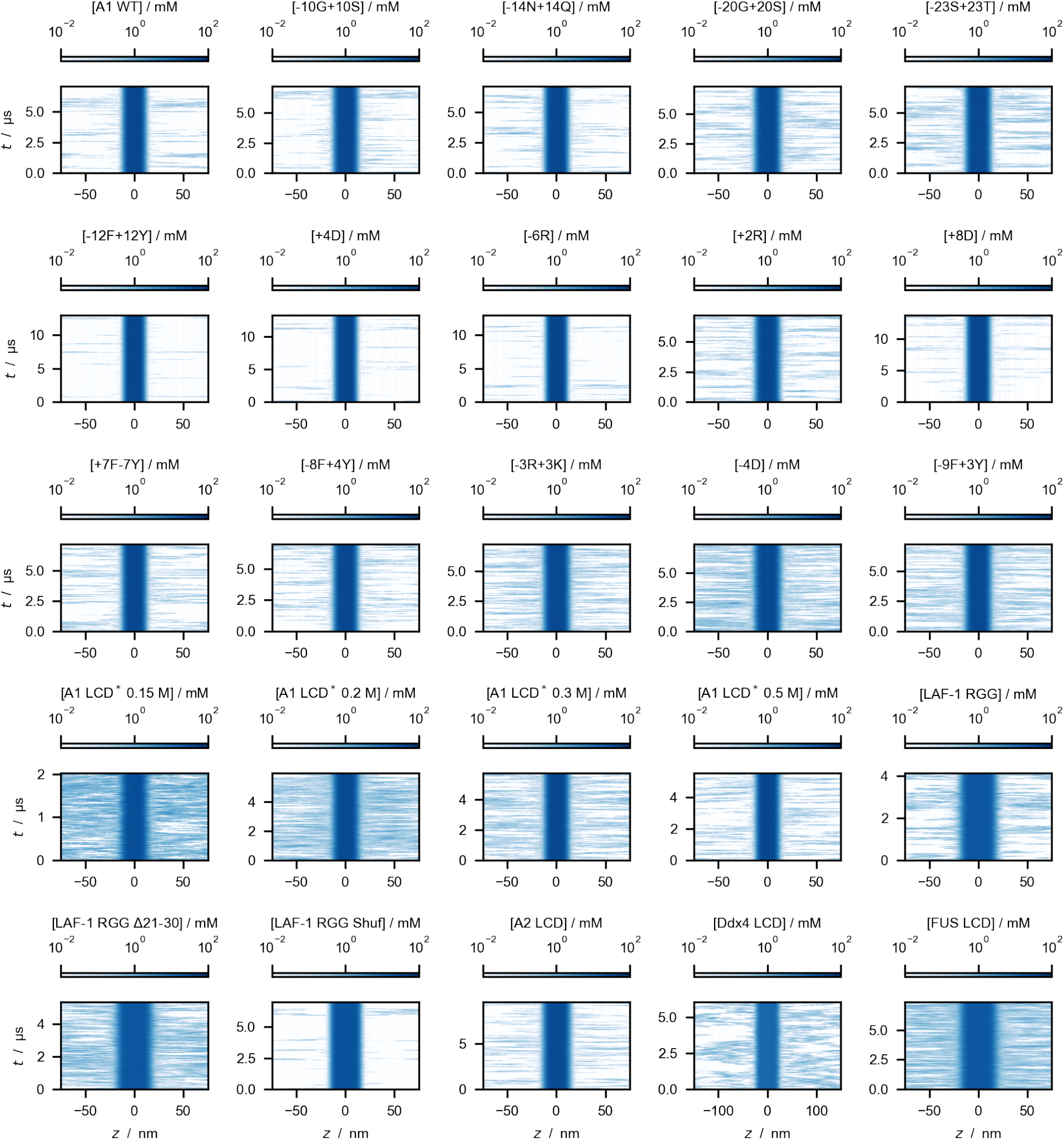
Time evolution of protein density along the *z*-axis of the simulation cell, as obtained from direct-coexistence simulations performed with the CALVADOS 2 model and *r*_*c*_ = 2 nm.

**Figure S14:**
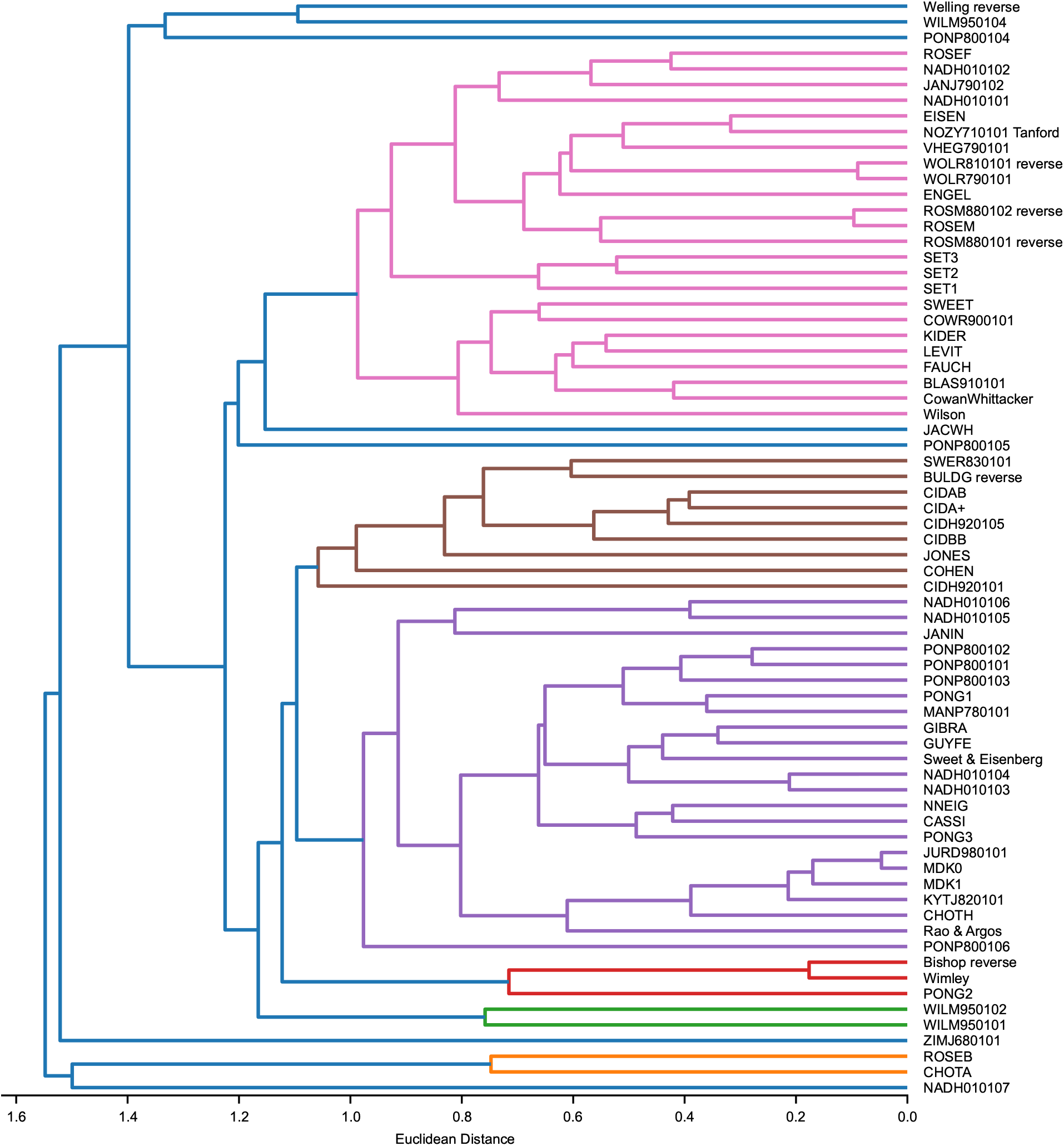
Hierarchical clustering dendrogram of 70 min-max normalized hydrophobicity scales selected from the set by Simm et al. [51]. Agglomerative clustering is performed using Euclidean distances and the average linkage method as implemented in the Python scikit-learn package [52].

**Figure S15:**
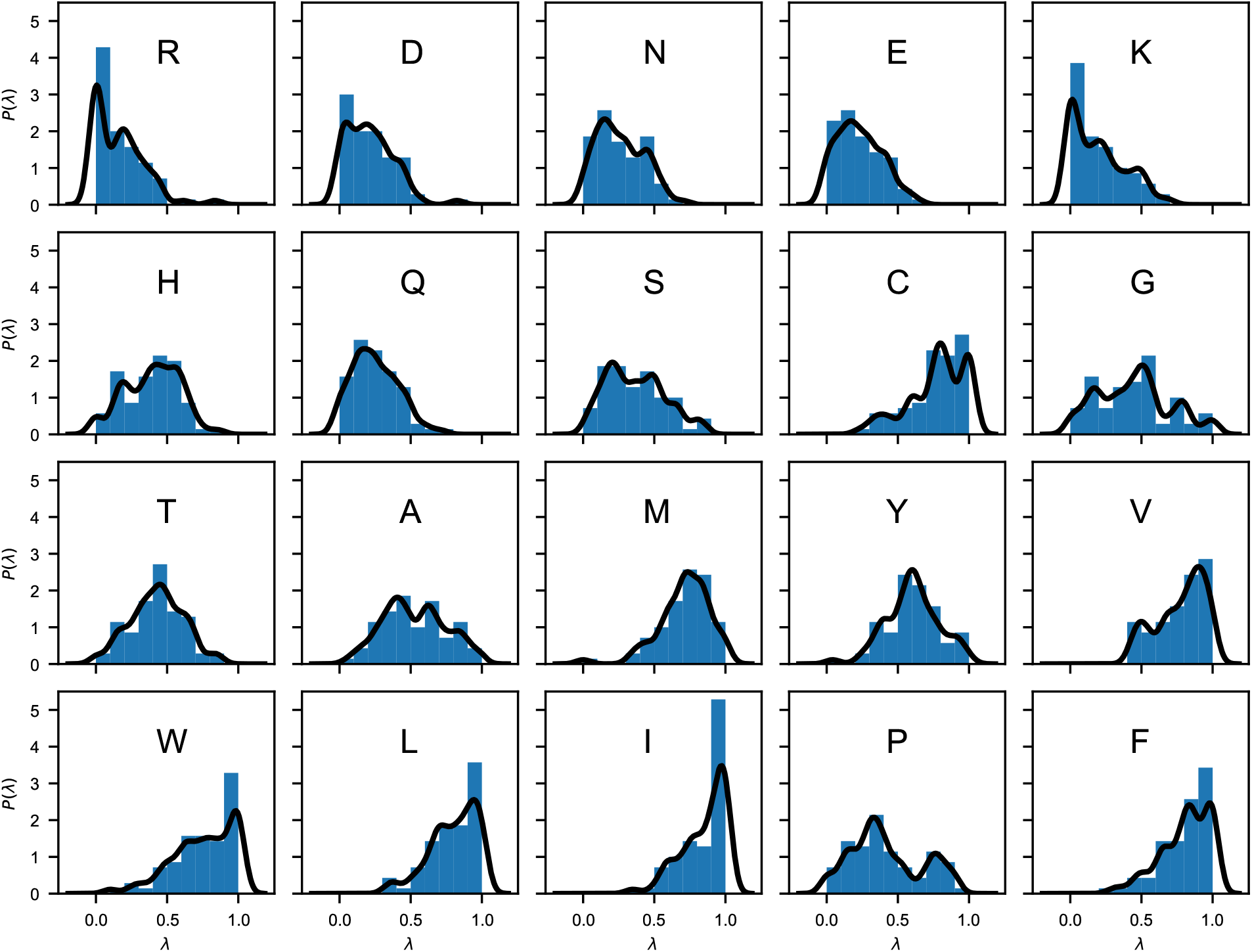
Probability distributions of the “stickiness” parameters, *P* (*λ*), obtained from 70 min-max normalized hydrophobicity scales selected from the set by Simm et al. [51]. Blue bars are histograms with bin width of 0.1. Black lines are obtained as 1D projections of the multivariate kernel density estimation implemented in scikit-learn [52], using a Gaussian kernel with bandwidth of 0.05.

**Figure S16:**
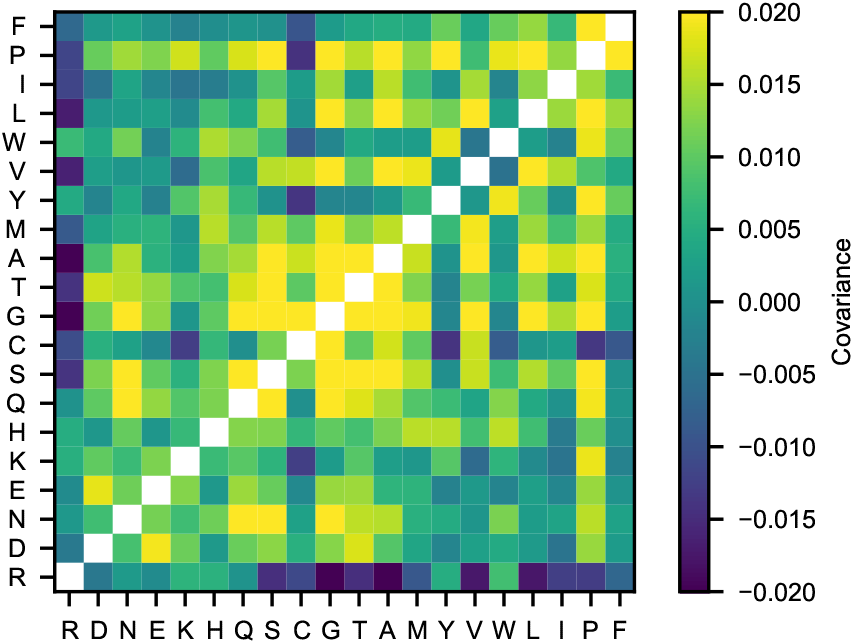
Covariance matrix of the 70 min-max normalized hydrophobicity scales selected from the set by Simm et al. [51]. The upper triangle of the matrix shows the covariance calculated directly from the 70 min-max normalized hydrophobicity scales whereas the lower triangle of the matrix shows the covariance calculated from the multivariate kernel density estimation averaging over the 70 min-max normalized hydrophobicity scales.

**Table S1:**
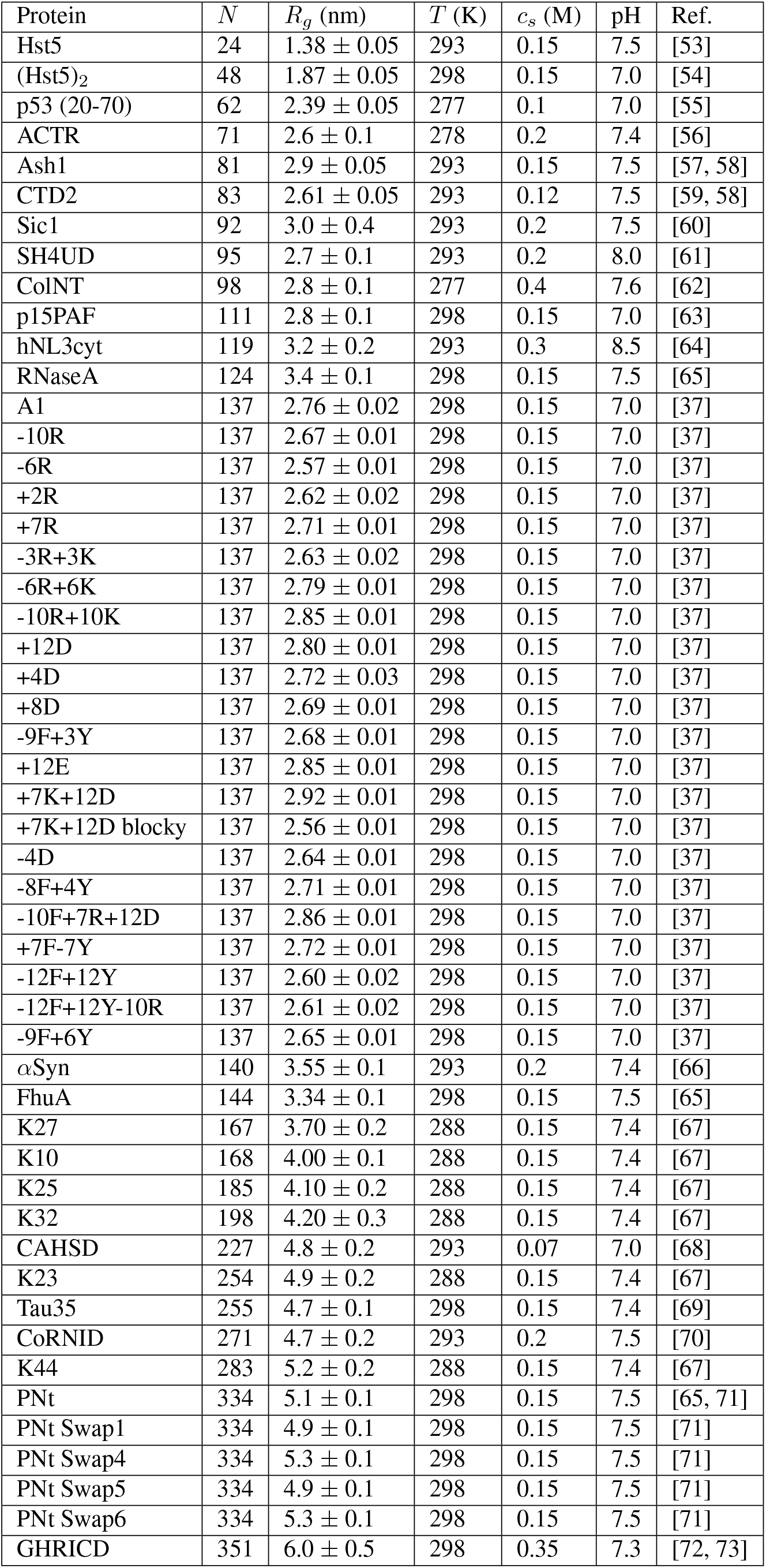
Solution conditions and experimental radii of gyration of proteins included in the training set for the Bayesian parameter-learning procedure.

| Protein | $N$ | $R_g$ (nm) | $T$ (K) | $c_s$ (M) | pH | Ref. |
| --- | --- | --- | --- | --- | --- | --- |
| Hst5 | 24 | $1.38 \pm 0.05$ | 293 | 0.15 | 7.5 | [53] |
| (Hst5)<sub>2</sub> | 48 | $1.87 \pm 0.05$ | 298 | 0.15 | 7.0 | [54] |
| p53 (20-70) | 62 | $2.39 \pm 0.05$ | 277 | 0.1 | 7.0 | [55] |
| ACTR | 71 | $2.6 \pm 0.1$ | 278 | 0.2 | 7.4 | [56] |
| Ash1 | 81 | $2.9 \pm 0.05$ | 293 | 0.15 | 7.5 | [57, 58] |
| CTD2 | 83 | $2.61 \pm 0.05$ | 293 | 0.12 | 7.5 | [59, 58] |
| Sic1 | 92 | $3.0 \pm 0.4$ | 293 | 0.2 | 7.5 | [60] |
| SH4UD | 95 | $2.7 \pm 0.1$ | 293 | 0.2 | 8.0 | [61] |
| ColNT | 98 | $2.8 \pm 0.1$ | 277 | 0.4 | 7.6 | [62] |
| p15PAF | 111 | $2.8 \pm 0.1$ | 298 | 0.15 | 7.0 | [63] |
| hNL3cyt | 119 | $3.2 \pm 0.2$ | 293 | 0.3 | 8.5 | [64] |
| RNaseA | 124 | $3.4 \pm 0.1$ | 298 | 0.15 | 7.5 | [65] |
| A1 | 137 | $2.76 \pm 0.02$ | 298 | 0.15 | 7.0 | [37] |
| -10R | 137 | $2.67 \pm 0.01$ | 298 | 0.15 | 7.0 | [37] |
| -6R | 137 | $2.57 \pm 0.01$ | 298 | 0.15 | 7.0 | [37] |
| +2R | 137 | $2.62 \pm 0.02$ | 298 | 0.15 | 7.0 | [37] |
| +7R | 137 | $2.71 \pm 0.01$ | 298 | 0.15 | 7.0 | [37] |
| -3R+3K | 137 | $2.63 \pm 0.02$ | 298 | 0.15 | 7.0 | [37] |
| -6R+6K | 137 | $2.79 \pm 0.01$ | 298 | 0.15 | 7.0 | [37] |
| -10R+10K | 137 | $2.85 \pm 0.01$ | 298 | 0.15 | 7.0 | [37] |
| +12D | 137 | $2.80 \pm 0.01$ | 298 | 0.15 | 7.0 | [37] |
| +4D | 137 | $2.72 \pm 0.03$ | 298 | 0.15 | 7.0 | [37] |
| +8D | 137 | $2.69 \pm 0.01$ | 298 | 0.15 | 7.0 | [37] |
| -9F+3Y | 137 | $2.68 \pm 0.01$ | 298 | 0.15 | 7.0 | [37] |
| +12E | 137 | $2.85 \pm 0.01$ | 298 | 0.15 | 7.0 | [37] |
| +7K+12D | 137 | $2.92 \pm 0.01$ | 298 | 0.15 | 7.0 | [37] |
| +7K+12D blocky | 137 | $2.56 \pm 0.01$ | 298 | 0.15 | 7.0 | [37] |
| -4D | 137 | $2.64 \pm 0.01$ | 298 | 0.15 | 7.0 | [37] |
| -8F+4Y | 137 | $2.71 \pm 0.01$ | 298 | 0.15 | 7.0 | [37] |
| -10F+7R+12D | 137 | $2.86 \pm 0.01$ | 298 | 0.15 | 7.0 | [37] |
| +7F-7Y | 137 | $2.72 \pm 0.01$ | 298 | 0.15 | 7.0 | [37] |
| -12F+12Y | 137 | $2.60 \pm 0.02$ | 298 | 0.15 | 7.0 | [37] |
| -12F+12Y-10R | 137 | $2.61 \pm 0.02$ | 298 | 0.15 | 7.0 | [37] |
| -9F+6Y | 137 | $2.65 \pm 0.01$ | 298 | 0.15 | 7.0 | [37] |
| αSyn | 140 | $3.55 \pm 0.1$ | 293 | 0.2 | 7.4 | [66] |
| FhuA | 144 | $3.34 \pm 0.1$ | 298 | 0.15 | 7.5 | [65] |
| K27 | 167 | $3.70 \pm 0.2$ | 288 | 0.15 | 7.4 | [67] |
| K10 | 168 | $4.00 \pm 0.1$ | 288 | 0.15 | 7.4 | [67] |
| K25 | 185 | $4.10 \pm 0.2$ | 288 | 0.15 | 7.4 | [67] |
| K32 | 198 | $4.20 \pm 0.3$ | 288 | 0.15 | 7.4 | [67] |
| CAHSD | 227 | $4.8 \pm 0.2$ | 293 | 0.07 | 7.0 | [68] |
| K23 | 254 | $4.9 \pm 0.2$ | 288 | 0.15 | 7.4 | [67] |
| Tau35 | 255 | $4.7 \pm 0.1$ | 298 | 0.15 | 7.4 | [69] |
| CoRNID | 271 | $4.7 \pm 0.2$ | 293 | 0.2 | 7.5 | [70] |
| K44 | 283 | $5.2 \pm 0.2$ | 288 | 0.15 | 7.4 | [67] |
| PNt | 334 | $5.1 \pm 0.1$ | 298 | 0.15 | 7.5 | [65, 71] |
| PNt Swap1 | 334 | $4.9 \pm 0.1$ | 298 | 0.15 | 7.5 | [71] |
| PNt Swap4 | 334 | $5.3 \pm 0.1$ | 298 | 0.15 | 7.5 | [71] |
| PNt Swap5 | 334 | $4.9 \pm 0.1$ | 298 | 0.15 | 7.5 | [71] |
| PNt Swap6 | 334 | $5.3 \pm 0.1$ | 298 | 0.15 | 7.5 | [71] |
| GHRICD | 351 | $6.0 \pm 0.5$ | 298 | 0.35 | 7.3 | [72, 73] |

**Table S2:** Solution conditions and experimental radii of gyration of proteins simulated in this study but not included in the training set for the Bayesian parameter-learning procedure.

| Protein | $N$ | $R_g$ (nm) | $T$ (K) | $c_s$ (M) | pH | Ref. |
| --- | --- | --- | --- | --- | --- | --- |
| DSS1 | 71 | $2.5 \pm 0.1$ | 288 | 0.17 | 7.4 | [73] |
| p27Cv14 | 107 | $2.936 \pm 0.13$ | 293 | 0.095 | 7.2 | [42] |
| p27Cv15 | 107 | $2.915 \pm 0.10$ | 293 | 0.095 | 7.2 | [42] |
| p27Cv31 | 107 | $2.81 \pm 0.18$ | 293 | 0.095 | 7.2 | [42] |
| p27Cv44 | 107 | $2.492 \pm 0.13$ | 293 | 0.095 | 7.2 | [42] |
| p27Cv56 | 107 | $2.328 \pm 0.10$ | 293 | 0.095 | 7.2 | [42] |
| p27Cv78 | 107 | $2.211 \pm 0.03$ | 293 | 0.095 | 7.2 | [42] |
| PTMA | 111 | $3.7 \pm 0.2$ | 288 | 0.16 | 7.4 | [73] |
| NHE6cmdd | 116 | $3.2 \pm 0.2$ | 288 | 0.17 | 7.4 | [73] |
| A1 LCD* | 131 | $2.645 \pm 0.02$ | 293 | 0.05 | 7.5 | [41] |
| A1 LCD* | 131 | $2.65 \pm 0.02$ | 293 | 0.15 | 7.5 | [41] |
| A1 LCD* | 131 | $2.62 \pm 0.02$ | 293 | 0.3 | 7.5 | [41] |
| A1 LCD* | 131 | $2.528 \pm 0.02$ | 293 | 0.5 | 7.5 | [41] |
| ANAC046 | 167 | $3.6 \pm 0.3$ | 298 | 0.14 | 7.0 | [73] |
| Tau 2N3R | 410 | $6.3 \pm 0.3$ | 298 | 0.15 | 7.4 | [69] |
| Tau 2N4R | 441 | $6.7 \pm 0.3$ | 298 | 0.15 | 7.4 | [69] |

**Table S3:**
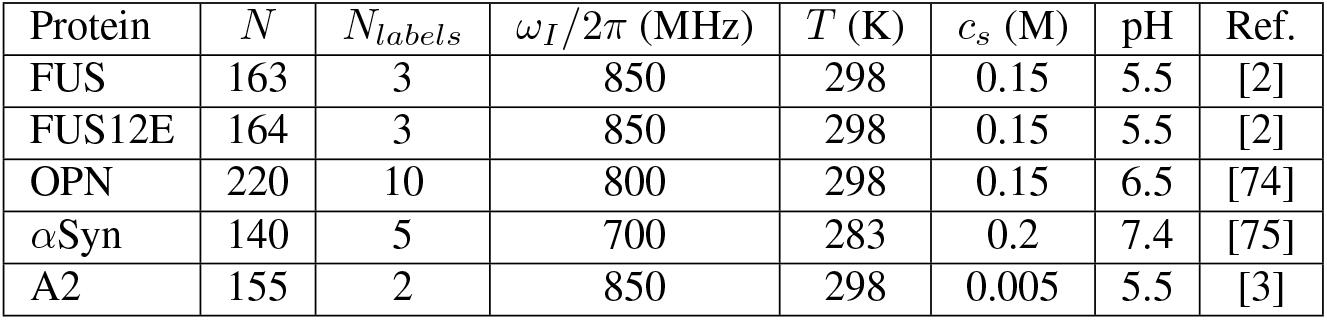
Experimental conditions for the intramolecular PRE data included in the training set.

**Table S4:** Conditions used for the direct-coexistence simulations performed in this study and references to the experimental data. Shaded rows highlight systems which are not included in the correlation plot of Fig. S7*B* because of lack of experimental *c*_*sat*_ values.

| Protein | N | $c_s$ (mM) | pH | Ref. | T (K) | | |
| --- | --- | --- | --- | --- | --- | --- | --- |
|  |  |  |  |  | 4 nm | 2 nm | Fig. 1C |
| 6His-TEV-Lge1<sub>1-80</sub>-StrepII WT | 114 | 500 | 7.5 | [43] | - | 293 | - |
| 6His-TEV-Lge1<sub>1-80</sub>-StrepII -11R+11K | 114 | 500 | 7.5 | [43] | - | 293 | - |
| 6His-TEV-Lge1<sub>1-80</sub>-StrepII -14Y+14A | 114 | 500 | 7.5 | [43] | - | 293 | - |
| A1 LCD WT | 137 | 150 | 7.0 | [4, 37] | 310 & 323 | 277 & 293 | 310 |
| A1 LCD +7F-7Y | 137 | 150 | 7.0 | [37] | 310 & 323 | 277 & 293 | - |
| A1 LCD -12F+12Y | 137 | 150 | 7.0 | [37] | 310 & 323 | 277 & 293 | - |
| A1 LCD -23S+23T | 137 | 150 | 7.0 | [37] | 310 & 323 | 277 & 293 | - |
| A1 LCD -14N+14Q | 137 | 150 | 7.0 | [37] | 310 & 323 | 277 & 293 | - |
| A1 LCD -10G+10S | 137 | 150 | 7.0 | [37] | 310 & 323 | 277 & 293 | - |
| A1 LCD -20G+20S | 137 | 150 | 7.0 | [37] | 310 & 323 | 277 & 293 | - |
| A1 LCD -30G+30S | 137 | 150 | 7.0 | [37] | 323 | 293 | - |
| A1 LCD +23G-23S | 137 | 150 | 7.0 | [37] | 323 | 293 | - |
| A1 LCD +23G-23S+7F-7Y | 137 | 150 | 7.0 | [37] | 323 | 293 | - |
| A1 LCD +23G-23S-12F+12Y | 137 | 150 | 7.0 | [37] | 323 | 293 | - |
| A1 LCD -9F+3Y | 137 | 150 | 7.0 | [37] | 310 | 277 | - |
| A1 LCD -8F+4Y | 137 | 150 | 7.0 | [37] | 310 | 277 | - |
| A1 LCD -3R+3K | 137 | 150 | 7.0 | [37] | 310 | 277 | - |
| A1 LCD -6R | 137 | 150 | 7.0 | [37] | 310 | 277 | - |
| A1 LCD -4D | 137 | 150 | 7.0 | [37] | 310 | 277 | - |
| A1 LCD +4D | 137 | 150 | 7.0 | [37] | 310 | 277 | - |
| A1 LCD +8D | 137 | 150 | 7.0 | [37] | 310 | 277 | - |
| A1 LCD +2R | 137 | 150 | 7.0 | [37] | 310 | 277 | - |
| A1 LCD* WT | 131 | 150 | 7.0 | [76] | 323 | 293 | - |
| A1 LCD* WT | 131 | 200 | 7.0 | [76] | 323 | 293 | - |
| A1 LCD* WT | 131 | 300 | 7.0 | [76] | 323 | 293 | - |
| A1 LCD* WT | 131 | 500 | 7.0 | [76] | 323 | 293 | - |
| LAF-1 RGG Domain | 176 | 150 | 7.5 | [77] | 323 | 293 | 293 |
| LAF-1 RGG Domain Shuffled | 176 | 150 | 7.5 | [77] | 323 | 293 | 323 |
| LAF-1 RGG Domain Δ21-30 | 166 | 150 | 7.5 | [77] | 323 | 293 | - |
| A2 LCD | 155 | 10 | 5.5 | [78] | - | 297 | - |
| FUS LCD | 163 | 150 | 7.4 | [79] | - | 297 | - |
| Ddx4 LCD | 236 | 130 | 6.5 | [80] | - | 297 | - |
| Human Full-Length Tau (2N4R) | 441 | 70 | 7.4 | - | - | - | 277 |

